# Rapamycin microparticles induce autophagy, prevent senescence and are effective in treatment of Osteoarthritis

**DOI:** 10.1101/2021.07.20.453073

**Authors:** Kaamini M. Dhanabalan, Ameya A. Dravid, Smriti Agarwal, Ramanath K. Sharath, Ashok K. Padmanabhan, Rachit Agarwal

**Affiliations:** Centre for BioSystems Science and Engineering, Indian Institute of Science, Bengaluru, India 560012.; Department of Orthopaedics, M.S. Ramaiah Medical College, Bengaluru, India 560054.

## Abstract

Trauma to the knee joint is associated with significant cartilage degeneration and erosion of subchondral bone, which eventually leads to osteoarthritis (OA), resulting in substantial morbidity and healthcare burden. With no disease-modifying drugs in clinics, the current standard of care focuses on symptomatic relief and viscosupplementation. Modulation of autophagy and targeting senescence pathways are emerging as potential treatment strategies. Rapamycin has shown promise in OA disease amelioration by autophagy upregulation, yet its clinical use is hindered by difficulties in achieving therapeutic concentrations, necessitating multiple weekly injections. Here, we have synthesized rapamycin - loaded poly (lactic-co-glycolic acid) microparticles (RMPs) that induced autophagy, prevented senescence and sustained sulphated glycosaminoglycans(sGAG) production in primary human articular chondrocytes from OA patients. RMPs were potent, nontoxic, and exhibited high retention time (up to 35 days) in mice joints. Intra-articular delivery of RMPs effectively mitigated cartilage damage and inflammation in surgery-induced OA when administered as a prophylactic or therapeutic regimen. Together, our studies demonstrate the feasibility of using RMPs as a potential clinically translatable therapy to prevent and treat post-traumatic osteoarthritis.

**Graphical Abstract:** 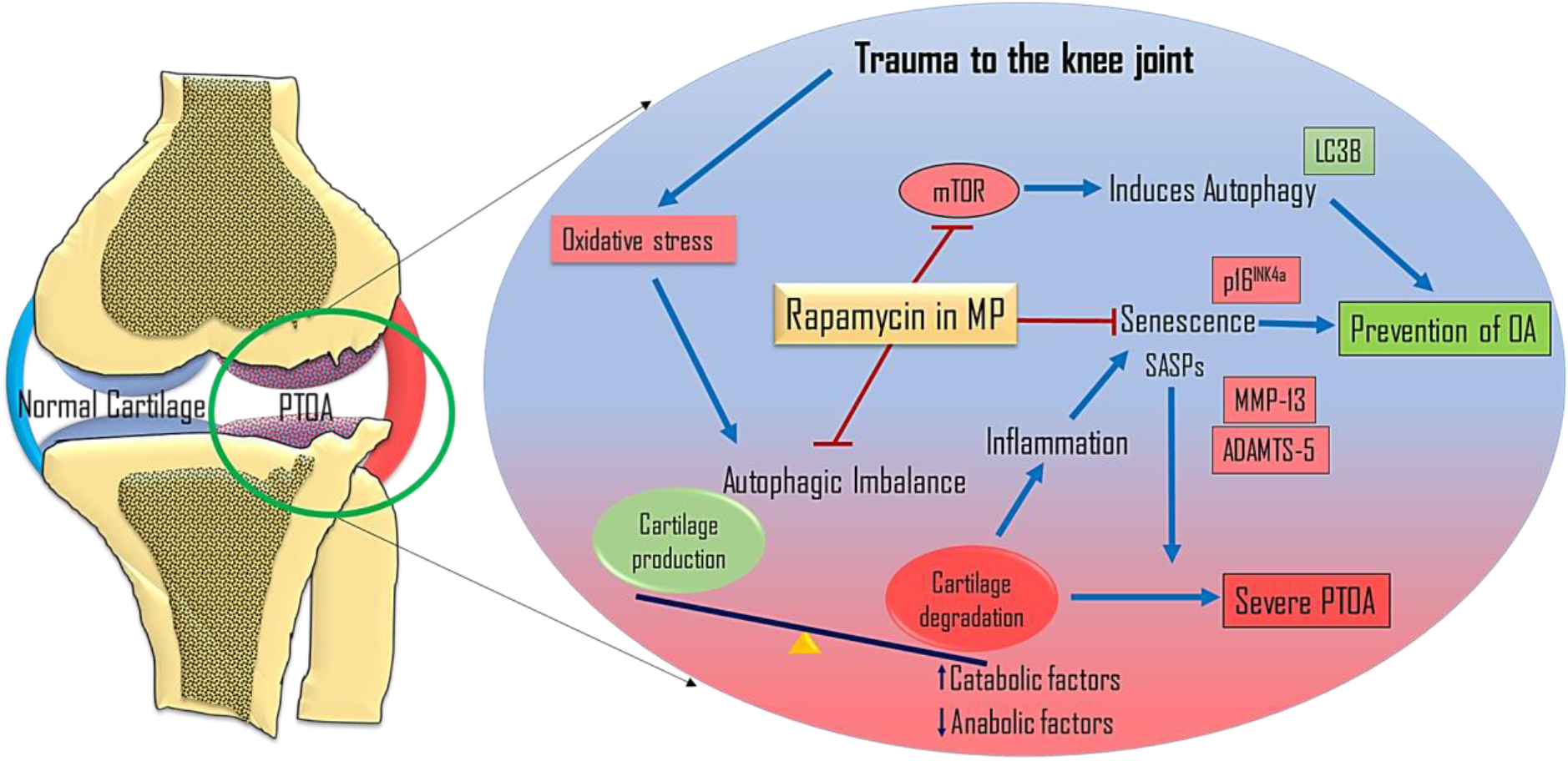

---

Osteoarthritis, a chronic progressive joint disorder, has affected nearly 500 million people worldwide and these numbers are projected to increase in the coming years which poses a huge societal burden and health care expense(*1*). As a progressive degenerative disease, cartilage damage can lead to synovitis, osteophyte formation, and hypertrophy of the entire joint capsule, ultimately leading to loss of function(*2*). Acute damage to the cartilage can occur with trauma to the joint such as vehicle accidents, falls, sports injuries, and military activities. Chronic damage occurs with aging, coupled with obesity, diabetes, and hormonal imbalances(*2*). The current treatment strategies in clinics, such as non-steroidal anti-inflammatory drugs, steroids, local viscosupplementation and physiotherapy, address only the symptoms like pain and inflammation(*3, 4*). There is a lack of any disease-modifying drug at the clinics that can prevent, halt, or reverse the disease progression(*5*).

Growing evidence points at deficient nutrient recycling mechanisms during stress conditions (mechanical, obesity, aging, injuries etc.) and an imbalance of the catabolic and anabolic pathways as the primary etiology of OA(*6–11*). Autophagy is a cellular homeostasis mechanism that maintains the balance between catabolic and anabolic pathways, and autophagic imbalance in chondrocytes is widely implicated in the onset and progress of OA(*12–16*). Failure in autophagy upregulation during these stress conditions, can make the cells apoptotic leading to diminished repair and remodelling capability in cartilage(*12, 13, 17–19*).

Chondrocytes under constant oxidative or genotoxic stress conditions can also turn senescent, leading to the secretion of inflammatory cytokines(*20–22*). The senescent chondrocytes secrete a host of inflammatory proteins such as IL-6, IL-8, and IL-17, together known as senescent associated secretory phenotype factors (SASP), which attracts immune cells leading to chronic inflammation(*23, 24*). Selective modulation of senescent cells has been one of the widely sought-after OA disease modifying strategies, where a wide variety of senolytic or senomorphic drugs are being evaluated, with a few of them undergoing clinical trials(*25–29*). Together, these evidences suggest that the autophagy activation and senescence modulation are critical for maintaining homeostasis of the articular cartilage and targeting these mechanisms can act as novel treatments to modify OA outcomes (*26, 27, 29–33*).

Rapamycin, a well-known immune modulator and an antibiotic used routinely in clinics for various diseases, has been shown to delay the progression of the OA in mice models(*30, 34*). Rapamycin induces autophagy via mTOR inhibition and pushes the cells from stress into the nutrient survival pathway and has shown to restore the health of the chondrocytes(*30, 34, 35*). Rapamycin also prevents senescence and stalls the production of SASP factors(*36–40*). It is hence, evident that rapamycin is a promising drug for OA treatment. However, free drug administration via intra-articular injection is challenging due to effective lymphatic clearance in joints and hence necessitates high dose administration and frequent injections. This can lead to toxicity, pain, joint infections and a considerable decrease in patient compliance.

Studies that have attempted to deliver OA disease-modifying agents such as inhibitors of catabolic enzymes and cytokine receptors via intra-articular injections have met with little clinical success(*41–43*). One of the main reasons for the lack of success of these potential disease-modifying agents is an insufficient therapeutic concentration for prolonged duration in the joint, owing to the efficient and rapid lymphatic clearance inside the knee joint(*44*). Senolytic drug, UBX0101, which was promising in treating OA in mice (*25*), recently failed to show efficacy after 12 week of a single intra-articular injection (maximum dose of 4 mg per patient) in phase two clinical trial (NCT04129944). It is hypothesized that the lack of efficacy of UBX0101 was due to its rapid clearance from the joints. Another phase I clinical trial using two injections (4 mg each) at week 0 and week 4 is currently underway (NCT04229225). Thus, it becomes evident that sufficient and prolonged therapeutic concentration inside the articular joints necessitate several frequent administrations, which significantly reduces patient compliance at clinics.

Intra-articular drug delivery systems using polymer-based slow-release formulation can serve as the key to bring many potent drugs into clinics(*45, 46*). Poly lactic-co-glycolic acid-based system (PLGA) is a robust and widely used clinical drug delivery system(*46, 47*). We had previously shown that PLGA based drug delivery systems have prolonged retention time in the murine knee joint(*38*). Therefore, we hypothesized that, intra-articular injections of rapamycin in PLGA based microparticles (RMPs) could prolong the drug’s residence time and could be used to treat OA.

Here, we report that the rapamycin PLGA microparticles induce autophagy and prevents senescence in primary human articular chondrocytes obtained from OA patients. The formulation sustained the production of sGAG production in stressed micromass cultures. The rapamycin particle formulation ameliorate surgery-induced OA in mice when administered as a prophylactic or a therapeutic regimen. To our knowledge, this is the first report of successful mice OA therapy using rapamycin in an injectable microparticle formulation and such strategies should be explored further for translation to humans.

## Results

### Chondrocytes from OA patients exhibited high senescence and diminished autophagy near OA lesions

To determine the basal autophagy and senescence burden of articular chondrocytes in end stage OA disease of humans, we examined the sections from explanted cartilage obtained from patients undergoing knee replacement surgery. Immunohistochemistry was used for staining respective markers of autophagy (LC3BII (microtubule associated protein 1B-Light chain 3)) and senescence (p16^INK4a^ (cyclin-dependent kinase inhibitor 2A)). Knee joints from OA patients undergoing total knee arthroplasty were obtained with appropriate consents and approvals. The tibial articulating surface was examined for gross morphology and the lesioned areas where, the subchondral bone erosion was evident were identified and labelled as OA-lesioned areas. Areas bearing healthy smooth cartilage surface morphology were identified as non-lesioned areas (**fig. S1**). The non-lesioned areas exhibited higher LC3B positive cells and comparatively fewer p16^INK4a^ positive cells suggesting the cartilage was healthy with active basal autophagy and fewer senescent cells (**fig. 1A** and **C**). However, the chondrocytes near the OA lesioned surface had lesser LC3B and high p16^INK4a^ positive cells, indicating that the cells near the OA lesioned cartilage exhibited high senescence and diminished autophagy (**fig. 1B** and **D**), which is in line with previously published results (*13, 48, 49*).

**Fig. 1.**
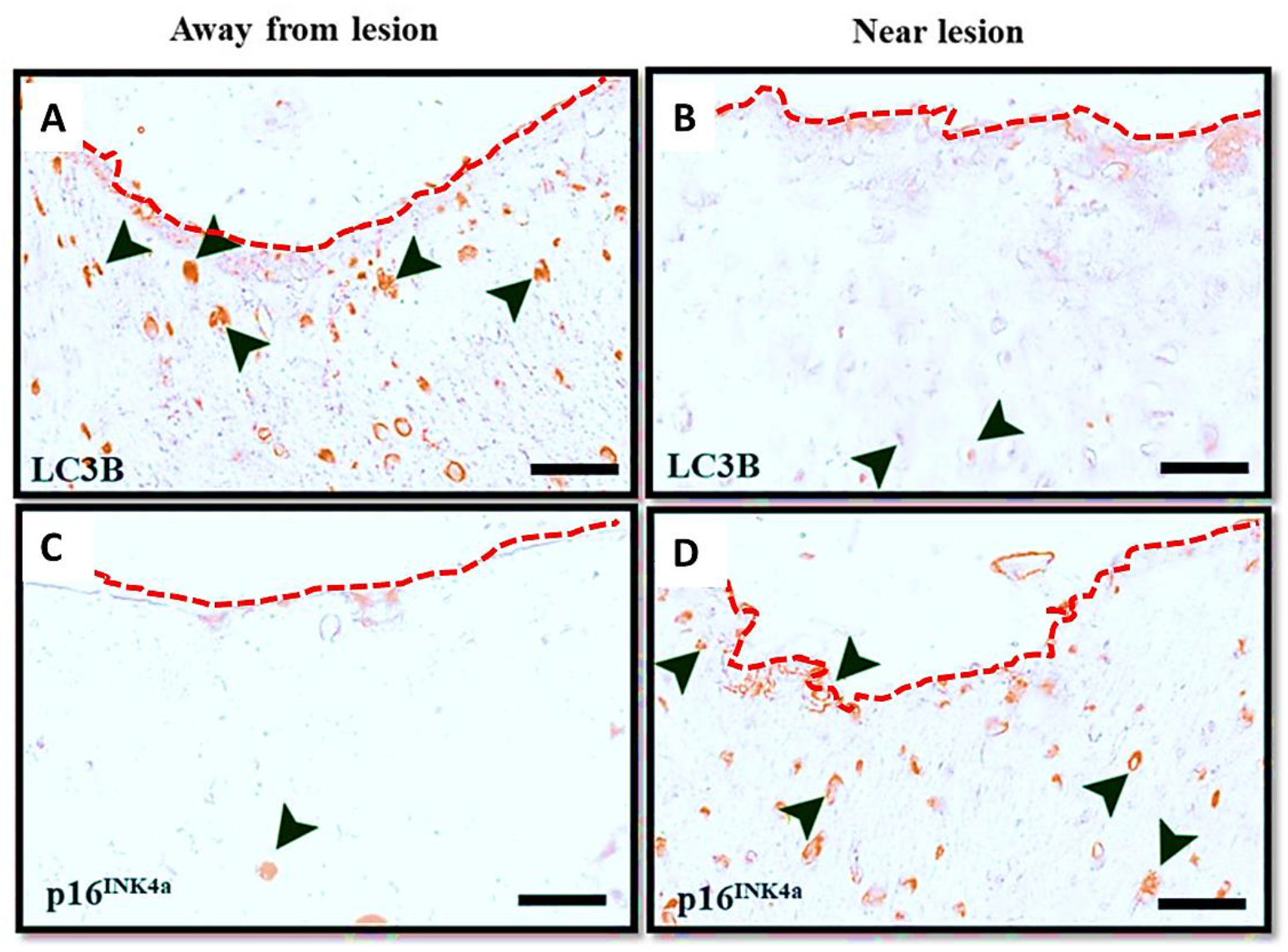
Chondrocytes from human osteoarthritic cartilage bear higher senescence burden and diminished autophagy near lesions. Immunohistochemical staining of LC3B in human osteoarthritic cartilage sections obtained from regions **(A)** away from lesion and **(B)** near OA lesion in knee joint cartilage. Immunohistochemical staining of p16^INK4a^ in human osteoarthritic cartilage sections obtained from regions **(C)** away from lesion and **(D)** near OA lesion in knee joint cartilage. Black arrows indicate the cells positive for the respective markers. The red dotted line represents the articular cartilage surface. The images shown are representative of data collected from three OA patients who underwent a total knee replacement. Scale bar, 25 μm.

### PLGA microparticles provide a tuneable platform to release rapamycin

Size plays an important role in prolonged joint retention time of PLGA microparticles. We had previously shown that particles in the size range of 1 μm exhibited high residence time in murine(*38*) and rodent(*50*) knee joints and hence we proceeded to use this size range for our current study. Dynamic light scattering analysis revealed an average diameter of 1039 ± 188 nm (**fig. 2A** and **Table S1**) and the zeta potential was −18 mV due to the presence of acid end capped PLGA polymer in the particles (**fig. 2B**).

**Fig. 2.**
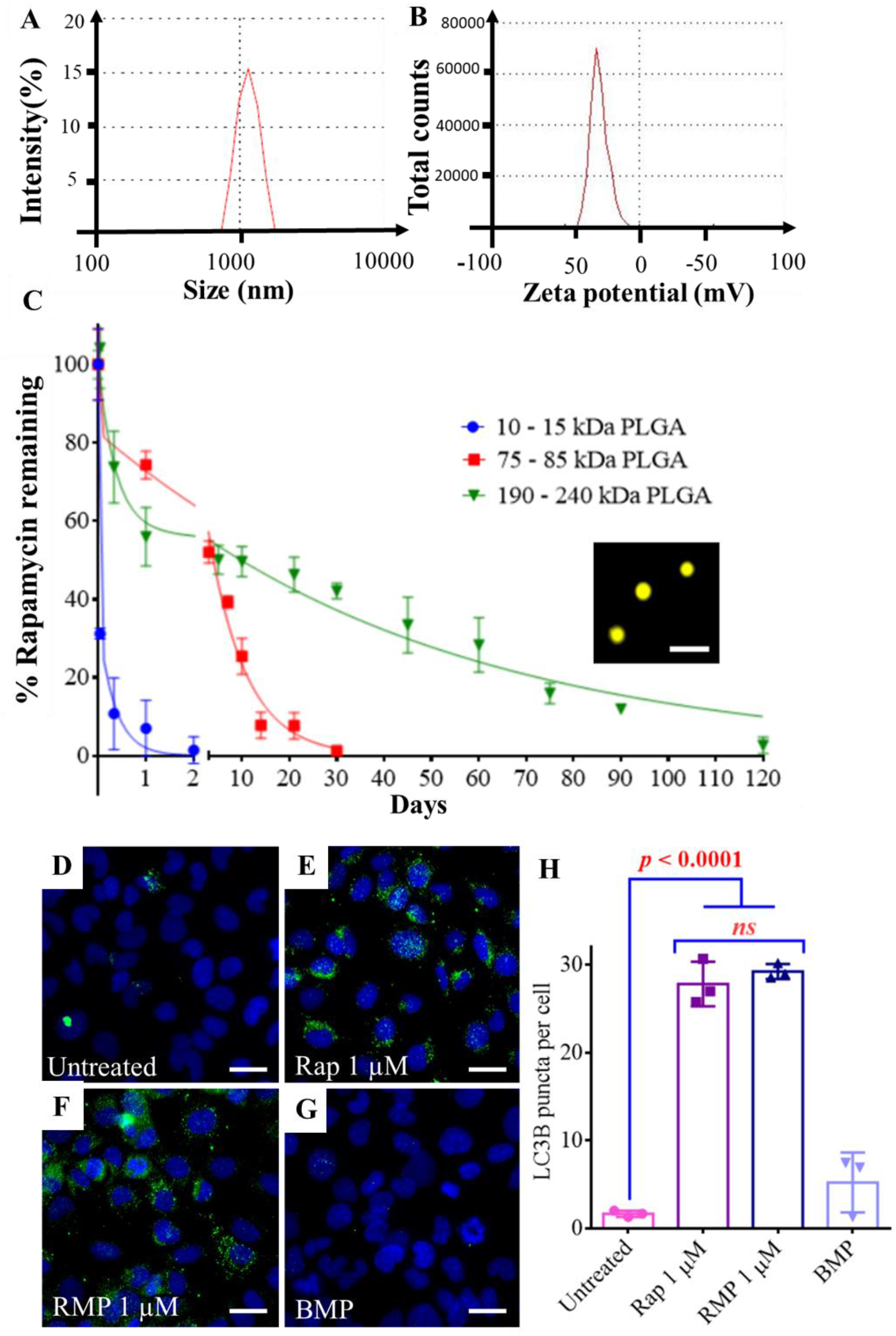
Rapamycin microparticles induce autophagy in primary human chondrocytes. **(A)** Size distribution of 10 - 15 kDa PLGA particles measured by Dynamic Light Scattering. **(B)** Zeta potential of 10 - 15 kDa PLGA particles. **(C)** *In vitro* release profiles of rapamycin from microparticles synthesized from different molecular weight PLGA polymers (n = 3 per group at each time point). Inset in **(C)** shows fluorescence microscopy image of Cy3 loaded PLGA particles (molecular weight 10 - 15 kDa); Scale bar, 3 μm. Fluorescence microscopy images of HACs stained with DAPI and LC3B after **(D) no treatment**, **(E)** free rapamycin (1 μM), **(F)** RMPs (1 μM rapamycin) and, **(G)** BMPs treatment. **(H)** Quantification of LC3B puncta per cell using ImageJ software (n = 3 per group). Plots were representative of data collected from three OA patients. Data in graphs represent the mean ± s.d. and *p* values were determined by one-way analysis of variance (ANOVA) and Tukey’s post hoc tests. BMP - Blank Microparticles, RMP - Rapamycin loaded Microparticles, HACs – Human Articular Chondrocytes. *ns* - non significant; Scale bar, 10 μm.

To assess the tuneability, microparticles of different molecular weights of PLGA were used to encapsulate rapamycin. The size and rapamycin encapsulation efficiency are listed in supplementary **Table S1**. It was observed that the PLGA MPs of molecular weight 10 - 15 kDa released the drug as quickly as in 48 h while PLGA MPs of molecular weight 75 - 85 kDa continued to release the drug up to 45 days. We varied the PLA: PGA ratio of the PLGA and found that the 190 - 240 kDa PLGA containing PLA: PGA (85:15) could extend the release of the drug for 120 days (**fig. 2C**). Thus, the particle platform was tuneable and allowed release of rapamycin in a controlled and sustained manner.

### Rapamycin MPs (RMPs) induce autophagy in primary human articular chondrocytes (HACs)

Rapamycin is a well-known autophagy inducer which acts by inhibiting mTOR signalling pathway(*51*). We had previously shown that rapamycin as a free drug as well as in MP formulation induced autophagy in the C28/I2 (human chondrocyte cell line) (**fig. S2A** to **E**)(*38*). Here, we found that in the primary HACs, both free rapamycin as well as RMPs successfully induced autophagy as visualized by the LC3B puncta in the cells.

Untreated HACs showed a low diffuse signal with very few puncta indicative of basal autophagy (Fig. 2D). In contrast, upon after treatment with free rapamycin (1 μM) or RMPs (equivalent rapamycin dose of 1 μM) (**fig. 2E** and **F**) for 48 h, LC3BII signal was observed as punctate signals. The results from RMPs were comparable to that of the free rapamycin treated groups (**fig. 2E**). The LC3B puncta per cell was higher in free rapamycin and RMPs treated cells compared to untreated cells. The untreated cells had an average of less than 2 puncta per cell indicative of basal autophagy, while the rapamycin and RMP treated groups exhibited around 25 puncta per cell indicative of upregulated autophagy upon rapamycin treatment. (**fig. 2H**). These findings suggest that RMPs were able to induce autophagy in the HACs.

### RMPs prevented senescence in primary human articular chondrocytes

Chondrocytes in the knee joint get exposed to various stress conditions such as DNA damage and increased ROS production(*52–54*). To simulate these stress conditions in the chondrocytes, we subjected them to oxidative stress by addition of H_2_O_2_ (200 μM). We also co-treated few of the experimental groups with free rapamycin and RMPs (1 μM) for 48 h and stained the cells with SA-β Gal (senescent marker) (**fig. 3A** to **D**). HACs when exposed to 200 μM of oxidative stress agent (H_2_O_2_), resulted in 30% of cells becoming senescent, while the co-treatment with 1 μM RMPs brought down the percentage of senescent cells to less than 10%, which was comparable to cells with no oxidative stress (**fig. 3E**).

**Fig. 3.**
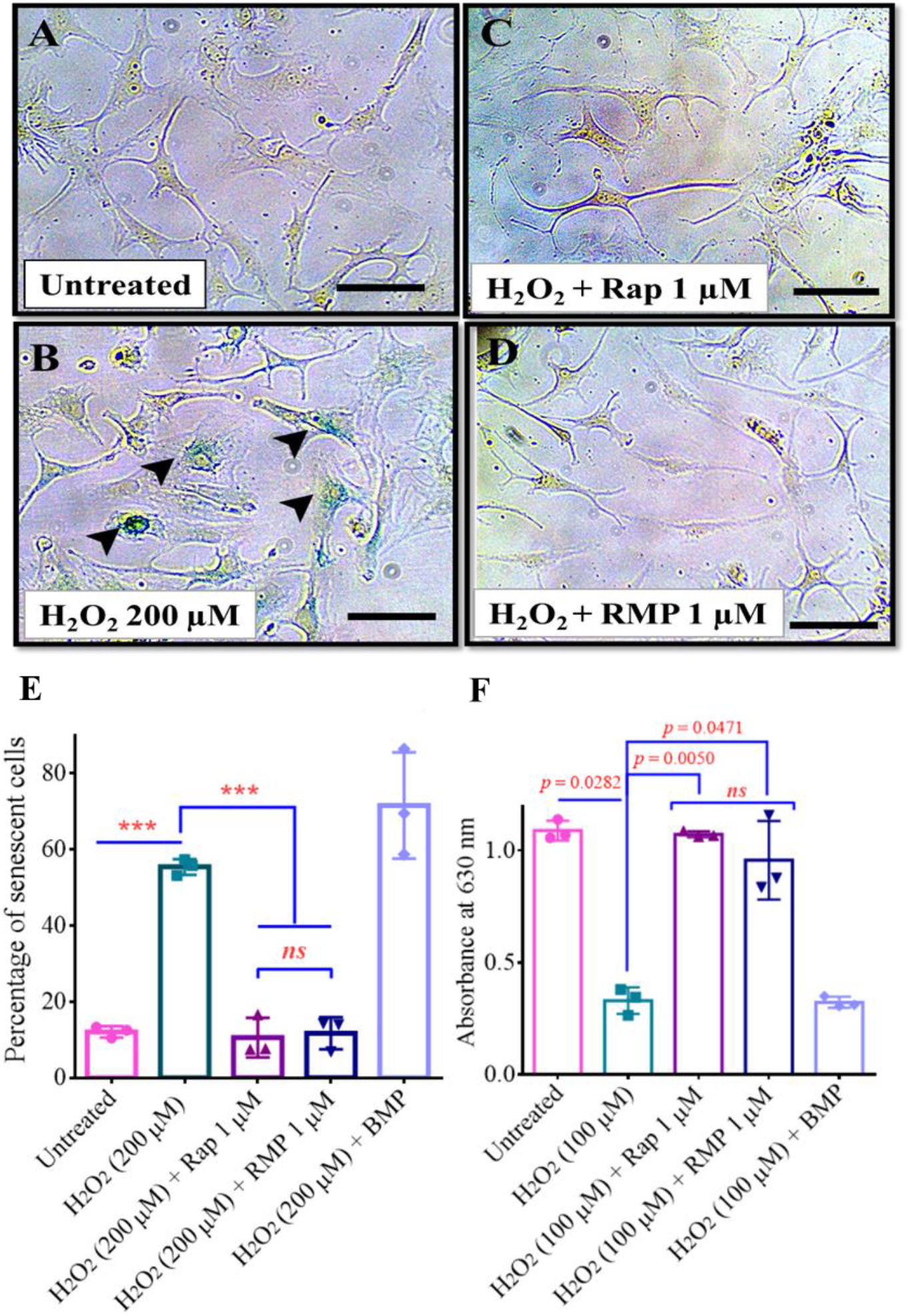
RMPs prevented senescence in primary HACs and sustained sGAG production in micromass cultures exposed to oxidative stress. SA-β Gal-stained images of primary HACs exposed to **(A) no treatment**, **(B)** oxidative (H_2_O_2_) stress, **(C)** oxidative (H_2_O_2_) stress with free rapamycin (1 μM) and, **(D)** oxidative (H_2_O_2_) stress with RMPs (1 μM). Scale bar, 40 μm. **(E)** Percent of senescent HACs after culturing in oxidative (H_2_O_2_) stress condition and different co-treatments (n = 3 per group) for 48 h. **(F)** sGAG production in stressed micromass cultures measured as absorbance of guanidine HCl extracted Alcian blue stain. Images and graphs were representatives of data collected from three OA patients. Data in graphs represent the mean ± s.d. and *p* values were determined by one-way analysis of variance (ANOVA) and Tukey’s post hoc tests. *p*-value < 0.05 was considered significant. HACs – Human Articular Chondrocytes, BMP - Blank Microparticles, RMP - Rapamycin loaded Microparticles. \*\*\*\**p*<0.0001, *ns* - non significant.

Similar results were also obtained with C28/I2 cell lines when cells were subjected to oxidative (H_2_O_2_, 200 μM) or genotoxic (BrdU, 200 μM) stress (**fig. S3-4**). Overall, these results indicate that rapamycin in MPs was potent and active post encapsulation and prevented senescence under different stress conditions in HACs and C28/I2 cell lines.

### RMPs helps to sustain sGAG production in stressed micromass cultures

When seeded at a high density along with the growth factor (TGF-β), chondrocytes form three-dimensional (3D) micromasses that exhibit high deposition of extracellular matrix components such as sulphated glycosaminoglycans (sGAG)(*55*). They serve as *in vitro* 3D culture models and are widely used in cartilage research to evaluate various stress conditions and treatment modalities. The micromasses derived from HACs were treated with oxidative stress using externally added H_2_O_2_, (100 μM) and co-treated with either free rapamycin or RMPs. We wanted to evaluate whether rapamycin or RMPs can sustain the production of sGAG in micromasses under oxidative stress conditions. In micromasses derived from HACs, H_2_O_2_ treatment resulted in nearly threefold lower sGAG production compared to the vehicle-treated groups while the free rapamycin or RMPs treatment rescued the sGAG production and was comparable to untreated groups (**fig. 3F**). Similar results were also obtained with micromasses formed from C28/I2 cells, when subjected to oxidative (H2O2, 100 μM) or genotoxic (BrdU, 600 μM) stress for 48 h (**fig. S5A** and **B**).

To evaluate the long term sGAG production by the stressed micromasses treated with rapamycin or RMPs, we treated micromass formed from C28/I2 cells, for 8 days with BrdU (600 μM)/ H_2_O_2_ (100 μM) along with co-treatment groups containing rapamycin or RMPs at 1 μM dose. In the BrdU (600 μM) treated group, the absorbance values dropped to almost six folds compared to untreated groups while the RMPs treated groups sustained the sGAG production on par with free rapamycin treated groups (**fig. S5B**). Likewise, sGAG production dropped to almost three folds with H_2_O_2_ treatment, and the rapamycin/RMPs treated group significantly increased the sGAG production at par with untreated groups (**fig. S5D**). In summary, these results suggest that rapamycin and RMPs were potent anabolic agents and sustained sGAG production in stressed chondrocytes for long durations under oxidative and genotoxic stress.

### PLGA MPs of high molecular weight had longer residence time in mice knee joints

In order to estimate the dosage and frequency of rapamycin to be administered for mice models of OA, we wanted to determine the residence time of rapamycin in the knee joint space. Since in our *in vitro* rapamycin release data, 75 -85 kDa PLGA MPs showed sustained release for more than a month (**fig. 2C**), we chose this formulation for all subsequent mice experiments. As it is difficult to assess the residence time of non-fluorescent drugs like rapamycin, we assessed the *in vitro* release rate of a fluorescent molecule, Cy7, which has similar molecular weight and size as rapamycin. The *in vitro* release profile of the RMPs and the Cy7 particles in 1x PBS (**fig. S6**) followed a similar trend and a two-phase decay curve using non-linear regression (least square method) was used to fit the release pattern of rapamycin and Cy7 dye from PLGA particles. Rapamycin and Cy7 dye followed a typical bi-phasic release from PLGA MPs, in line with the previously published literature(*38, 56, 57*).

Next, we injected Cy7 containing PLGA MPs intra-articularly and monitored the residence time inside the knee joints using an *in vivo* imaging system - Perkin Elmer IVIS^®^ Spectrum. The contralateral legs of mice received an equal concentration of Cy7 free dye (**fig. S7**). The joints receiving free dye only had 7% of the initial injected dose remaining on day 3, whereas the Cy7 MPs injected group showed fluorescent signal even on day 35 (**fig. S7C**). The joints did not show any gross signs of inflammation, infections, or difficulty in movements during the entire course of this study. These results were in line with our previously published work indicating that dyes encapsulated in 1 μm particles exhibit higher residence time inside mice and rodent knee joints compared to free dyes (*38, 50*). Taken together, these results suggest that the microparticle formulation was safe, did not elicit inflammation and exhibited prolonged joint residence time of up to 35 days compared to the free dye.

### RMPs prevented the progression of OA disease in murine post traumatic model of OA

To test our formulations on a functional model of OA, we utilized a widely used post-traumatic OA model in mice (destabilisation of the medial meniscus (DMM)). In this model, mice are subjected to destabilisation of the medial meniscus by nicking the medial menisco-tibial ligament and then left free to move in their cages which leads to development of severe OA by eight weeks(*58, 59*). To test the prophylactic capacity of the formulation, we administered Rapa, RMP and BMP intra-articularly in 2 doses, starting 1 week after the DMM (**fig. 4A**). To determine an optimal dose of rapamycin, we assessed two different doses (1.8 μg and 180 ng per joint) based on previously published literature(*30, 34*). The *in vitro* release curve of rapamycin from PLGA particles (75 – 85 kDa PLGA, **fig. 2C**) showed that 90% of the drug gets released in about 20 days, hence the injection time points (18-20 days apart) were chosen accordingly. In the BMPs and free rapamycin-treated groups, the cartilage surface appeared irregular, chondrocytes exhibited focal perichondral staining, and empty lacunae indicated chondrocyte death. RMPs at the dose of 1.8 μg per joint effectively prevented OA as the cartilage surface appeared smooth, and there was minimal loss of safranin stain. This suggests that the proteoglycan content was maintained, and there were fewer hypertrophic cells than vehicle-treated groups (**fig. 4B** and **C**). The OARSI scores of RMPs (1.8 μg dose) treated mice were four folds (*p* = 0.0008) and six folds (*p* ≤ 0.0001) lower than the free rapamycin (1.8 μg dose) treated and DMM mice groups respectively. The OARSI scores of BMPs treated mice were not different from DMM operated and free rapamycin treated groups and were statistically different from RMPs treated mice OARSI scores (*p* = 0.0003). The lower dose of 180 ng per joint, both as free drug as well as in MP formulation, failed to prevent the OA disease progression and hence, for all subsequent studies, we used a dose of 1.8 μg per joint.

**Fig. 4.**
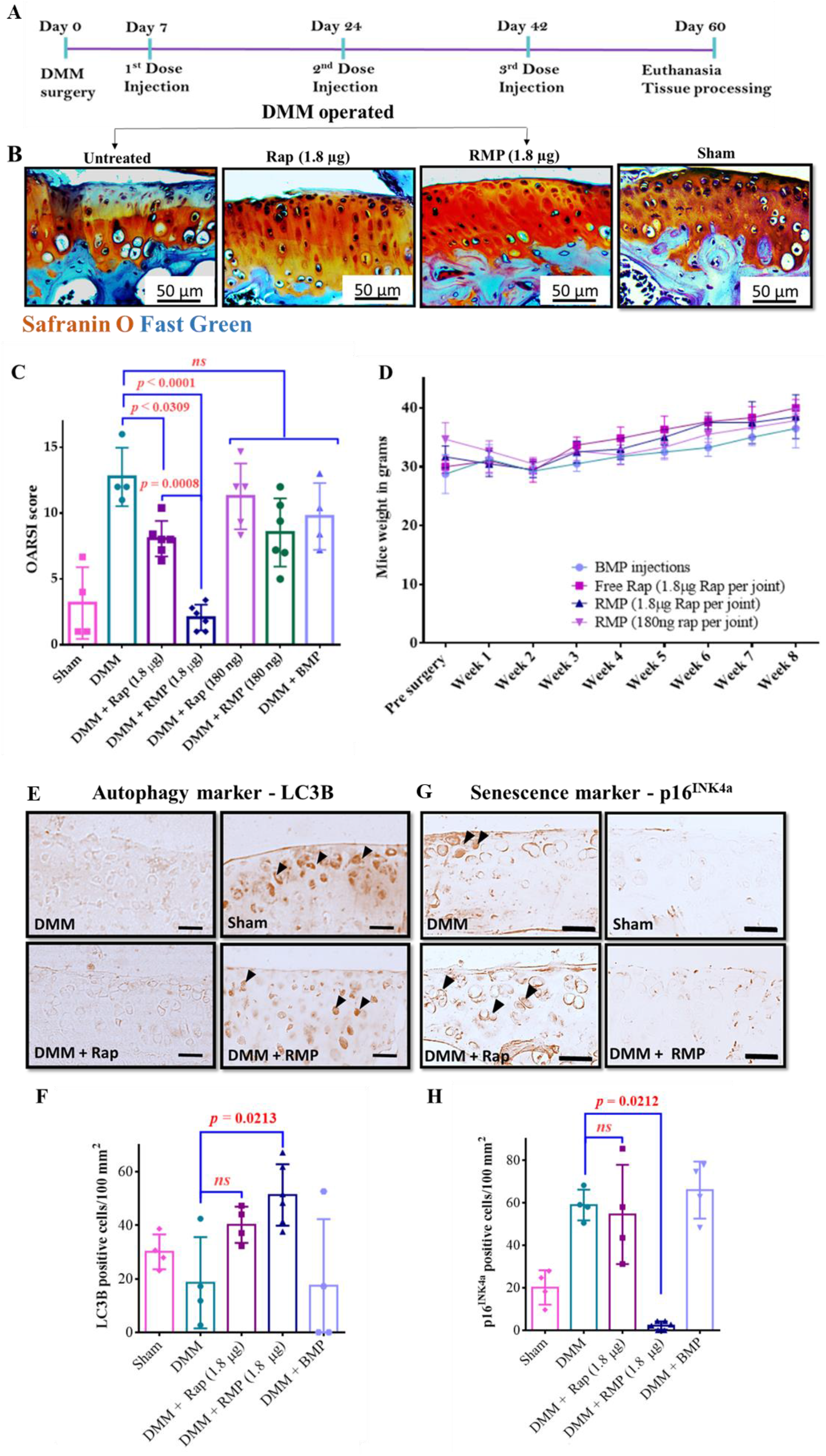
RMPs as a prophylactic regimen prevented the progression of OA in murine post-traumatic model of OA. **(A)** Schematic representation of prophylactic treatment timeline. **(B)** Representative Safranin-O and Fast Green–stained images of mice knee joints collected from different experimental groups. **(C)** Mouse OARSI scores showing the extent of injury in mice after receiving different treatments. **(D)** Plot showing bodyweight of mice with respect to time after receiving different treatments. **(E)** Representative images of LC3B IHC staining. **(F)** Quantification of LC3B positive cells per 100 mm^2^. **(G)** Representative images of p16^INK4a^ IHC staining. **(H)** Quantification of p16^INK4a^ positive cells per 100 mm^2^. The data in the graphs represent the mean ± s.d. and *p* values were determined using one-way analysis of variance (ANOVA) or Kruskal Wallis test and Tukey’s post hoc analysis. DMM (n = 4), sham (n = 4), rapamycin (high dose, n = 6; low dose n = 5), RMPs (n = 6), BMPs (n = 4). *ns* - non significant; Scale bar, 50 μm.

Immunohistochemical staining of the knee joints for inflammatory markers (ADAMTS-5 and MMP-13), autophagy marker (LC3B) and senescence marker (p16^INK4a^) were carried out on the joint sections. Consistent with our *in vitro* results, the articular cartilage surface of RMPs (1.8 μg) treated mice exhibited higher LC3B positive cells indicative of increased autophagy in the cells of the articular cartilage (**fig. 4E** and **F**). The free rapamycin treated group also exhibited LC3B positive cells, although they were not statistically different from the DMM and BMP treated group. The cells positive for p16^INK4a^, a senescence marker was more than 18-fold lower in the mice joints treated with 1.8 μg RMPs (**fig. 4F** and **H**) compared to DMM group (*p* = 0.0212). This could be attributed to the sustained and effective inhibition of the mTOR pathway by rapamycin which upregulates autophagy, thus preventing the cells from turning senescent(*30, 34, 35, 39*).

Autophagic imbalance in articular chondrocytes is associated with the up-regulation of matrix-degrading enzymes such as matrix metalloproteinases (MMPs) and a disintegrin and metalloprotease with thrombospondin motifs (ADAMTS). Of these proteases, MMP13 and ADAMTS-5 are critical players in OA progression as they degrade the extracellular matrix, and induce hypertrophy of the chondrocytes in the murine knee joints(*60–65*). We stained for MMP13 and ADAMTS-5 using IHC on the articular surface of different treatment groups. In the RMPs (1.8 μg) treated groups, the cartilage surface exhibited considerably lower MMP-13 and ADAMTS-5 positive cells than the DMM group (4-6-fold), and the number of positive cells was comparable to the sham mice group (**fig. S8A** to **E** and **S8F** to **J**). The DMM, free rapamycin-treated, and BMP-treated mice showed much higher MMP-13 and ADAMTS-5 staining compared to RMP-treated mice.

During the entire course of the treatment regimen, the mice were monitored for drug toxicity related weight loss. None of the animals in the different treatment groups exhibited any weight loss and showed steady weight gain (**fig. 4D**).

### Therapeutic regimen of RMPs reduces the severity of OA in murine post traumatic model of OA

Since most OA patients come to clinics only after experiencing symptoms, we wanted to evaluate whether our rapamycin formulations would be effective once the disease has substantially progressed. It has been shown in the literature that DMM operated mice start exhibiting significant early-stage OA hallmarks, such as proteoglycan loss, chondrocyte hypertrophy and cartilaginous osteophyte formation, as early as two weeks after surgery(*66*). Therefore, we administered the DMM operated mice groups with our formulation from three weeks post DMM surgery and further reduced the number of injections to two (21days apart) compared to three injections used in our prophylactic study (**fig. 5A**).

**Fig. 5.**
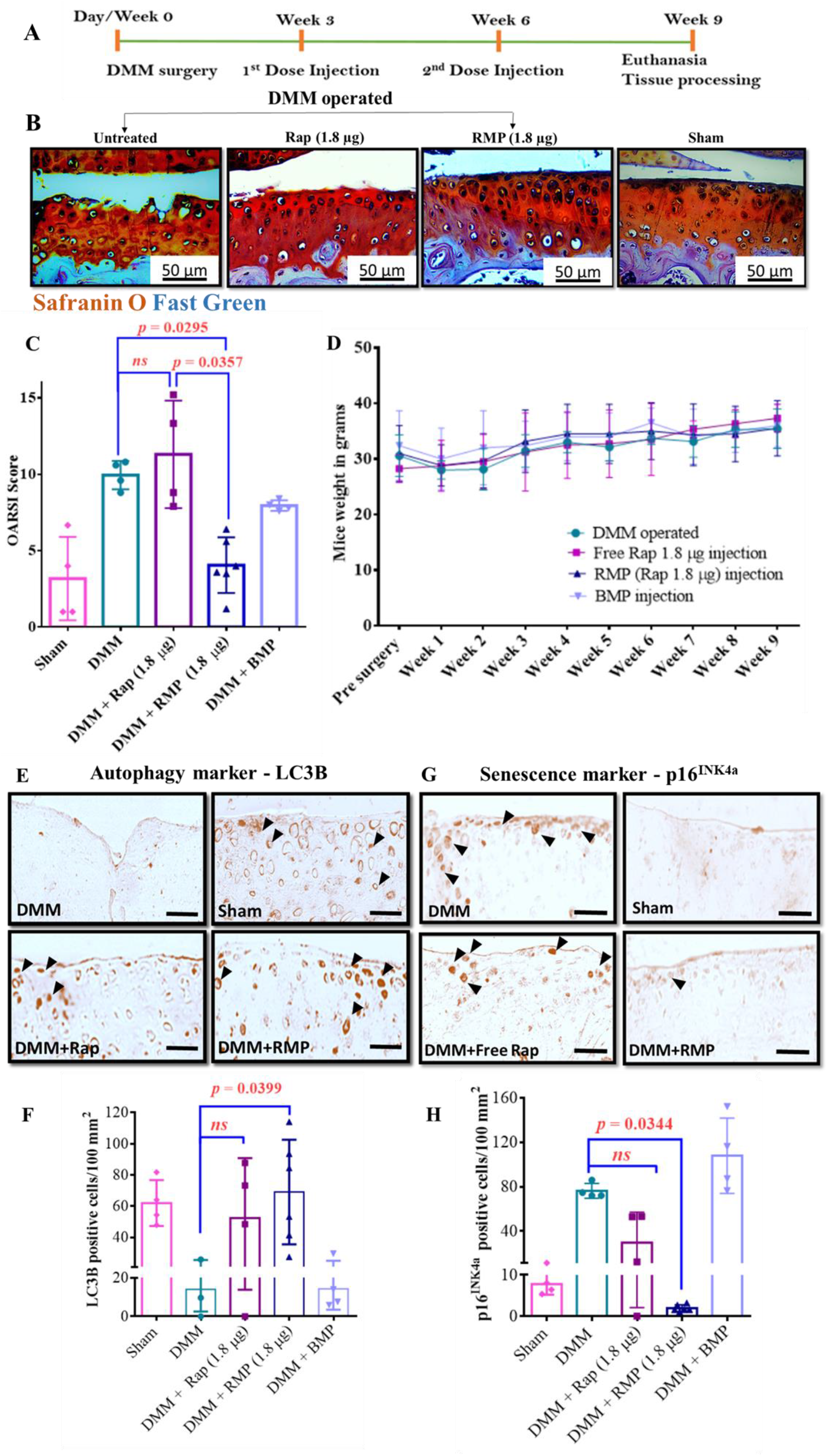
RMPs as a therapeutic regimen reduced the severity of OA in murine post traumatic model of OA. **(A)** Schematic representation of therapeutic treatment timeline. **(B)** Representative Safranin-O and Fast Green–stained images of mice knee joints collected from different experimental groups. **(C)** Mouse OARSI scores showing the extent of injury in mice after receiving different treatments. **(D)** Plot showing bodyweight of mice with respect to time after receiving different treatments. **(E)** Representative images of LC3B IHC staining. **(F)** Quantification of LC3B positive cells per 100 mm^2^. **(G)** Representative images of p16^INK4a^ IHC staining. **(H)** Quantification of p16^INK4a^ positive cells per 100 mm^2^. The data in the graphs represent the mean ± s.d. and *p* values were determined using one-way analysis of variance (ANOVA) or Kruskal Wallis test and Tukey’s post hoc analysis. DMM (n = 4), sham (n = 4), free rapamycin (n = 4), RMPs (n = 6), BMPs (n = 4). *ns* - non significant; Scale bar, 50μm.

Surgery-induced OA groups with no treatment, free rapamycin treatment, and BMP treatment showed reduced safranin O staining of proteoglycans accompanied by cartilage thinning, surface irregularities, hypertrophic chondrocytes, empty lacunae, and focal perichondral staining (**fig. 5B**). The OARSI scores of RMPs (1.8 μg) treated mice were 2.5-fold lower than the free rapamycin (1.8 μg) treated (*p* = 0.0357) and DMM only (*p* = 0.0295) mice OARSI scores. RMPs at dose of 1.8 μg per joint, effectively reduced these OA associated damages and improved the overall cartilage health.

Consistent with our *in vitro* results and prophylactic treatment regimen outcomes, the articular cartilage surface of RMPs (1.8 μg) treated mice exhibited a higher number of cells positive for LC3B, indicating active autophagy in the cells of the articular cartilage, while the free rapamycin and BMP treated groups exhibited fewer LC3B positive cells indicative of low basal autophagy (**fig. 5E** and **F**). The cells positive for p16^INK4a^ were more than 20-fold lower in the mice joints treated with 1.8 μg RMPs compared to free rapamycin or DMM only (*p* = 0.0344) groups (**fig. 5G** and **H**). The free rapamycin-treated group showed fewer senescent cells than the DMM and BMP-treated groups but were not statistically different. Consistent with our prophylactic studies, the markers of inflammation, MMP-13 and ADAMTS-5 positive cells on the articular surface of the DMM and BMP treated groups were higher in number indicative of cartilage degradation and inflammation. The cells positive for these inflammatory protease markers was more than 3-fold lower in RMP treated group compared to free rapamycin treated or DMM only groups (**fig. S9**). This is suggestive that sustained presence of rapamycin prevented the cells from secreting these proteases and sustained the ECM production, as evident by lower damage and uniform Safranin O staining (**fig. 5B** and **5C**, and **fig. S9A** to **E**).

Thus, the prophylactic and therapeutic regimens using RMPs in our surgically induced OA mice model effectively induced autophagy, prevented senescence, reduced the markers of inflammation in the joints, and paved the way for ameliorating OA in mice models.

## 4. Discussion

Several drugs and biologics like resveratrol, curcumin, torin1, NDGA, rapamycin, epigenetic modulators, senolytics and cytokine antagonists have been excellent in modifying OA pathology in animal models but have not entered clinics (*25, 31–34, 41, 67, 68*). This is attributed to the challenges in drug delivery, such as maintaining therapeutic concentrations and preventing rapid clearance of the drug from the targeted area(*5*). Furthermore, in several of these animal studies, multiple intra-articular injections were used, with some of them administering as high as two IA injections per week which are not clinically translatable. Therefore, the present investigation draws inspiration from two key concepts: the broad literature, which establishes rapamycin as a potent autophagy inducer and senomorphic drug(*30, 34, 35, 38, 69*), and engineering a sustained-release microparticle formulation for clinically acceptable and patient compliant therapy. Here we were able to show that an IA injection once every 3 weeks was sufficient to treat early OA in mice. This frequency of administration is routinely used in clinics as is the case with Hyalgan^®^ (3 cycles of weekly intra-articular injections for 5 weeks, NCT00669032), Enbrel^®^ (weekly intra-articular injections for 5 weeks, NCT02722772), Traumeel^®^/ Zeel^®^ (weekly intra-articular injections for 3 weeks, NCT01887678) and ongoing clinical trial with sprifermin (2 or 4 cycles of weekly intra-articular injections for 5 weeks, NCT01919164).

Recent research has shown that micro and nanoparticles can be used to synthesize sustained release systems and can be used to release the disease-modifying drugs and agents to ameliorate OA(*70, 71*). Utilizing such polymer-based sustained release systems could be the key to clinical translation of disease modifying drugs of osteoarthritis. Since PLGA based MPs are clinically used biomaterials with well-established safety profiles such as Zilretta® that is used for OA pain management, and as rapamycin is already clinically approved; rapamycin PLGA MPs can be rapidly translated to clinics.

For our animal experiments, we used microparticles synthesized from 75-85 kDa molecular weight PLGA which exhibited long residence time in (~35 days) in murine knee joints. It is possible that using PLGA particles made from higher molecular weight polymer may exhibit even longer residence time in the joints and further reduce the frequency of administration. In our *in vitro* rapamycin release profiles, we found that higher molecular weight PLGA (190-240 kDa) released rapamycin in 120 days compared to about 30 days for 75-85 kDa polymer. Such sustained release platforms should be tested and tuned with appropriate drug loading to allow continuous release of certain amount of drug and maintain therapeutic concentrations in the joints. Other DMOADs can also be co-delivered using this platform to explore synergy in OA treatment.

Drugs that exhibit multifaceted approaches such as autophagy activation, senescence prevention, and reducing inflammation in the milieu can potentially turn into promising therapies that are translatable(*72, 73*). Autophagy is a stress survival mechanism that gets activated as an adaptive process to different forms of metabolic stress, and the role of autophagy in osteoarthritis is widely studied(*8, 12, 15*). Cellular senescence is another growing field and is an important target for many diseases, including osteoarthritis(*48, 74–76*). In the mice model of OA, fewer cells were positive for LC3B (**fig. 4** and **5, E** and **F**) and a higher number of cells were positive for the senescence marker - p16^INK4a^ (**fig. 4** and **5, G** and **H**) compared to sham surgery group. In contrast, the RMPs treated groups upregulated autophagy (**fig. 4** and **5, E** and **F**) and reduced the expression of p16^INK4a^ (**fig. 4** and **5, G** and **H**) and chondrocyte hypertrophy (MMP13 and ADAMTS-5 expression). Similarly, *in vitro*, RMPs induced autophagy, successfully rescued oxidatively stressed primary human articular chondrocytes obtained from OA patients, from turning into senescent phenotypes, and sustained the production of sGAG (**fig. 2D-H** and **fig. 3**), which shows its clinical translation potential.

Many studies have highlighted the role of immune cells, such as T cells and B cells, in OA(*77, 78*). One of the emerging areas of investigation is evaluating the role of these immune cells, especially T cells in, OA progression. T helper(Th) cells such as Th1, Th9 and, Th17 are known to be present at a significantly higher number in OA joints(*79*). Studies show a positive feedback loop between Th17 cells, IL-17, and senescent synovial cells, with mTOR playing a significant role in upregulating the production of several cytokines, and clearing the Th17 cells can help in OA disease amelioration(*24, 80*). Rapamycin, being an immunosuppressive agent and mTOR signalling inhibitor, could directly act on immune cells and potentially reduce the secretion of many inflammatory cytokines such as IL-17, and thus can prevent protease-mediated destruction of cartilage(*35, 36, 80–82*). Further experiments are needed to determine whether rapamycin is involved in modulating Th cells and IL-17 production in OA. Since OA is predominantly a geriatric disease, and aged mice report a higher senescence burden and develop more severe OA after injury, additional evaluation into aged mice and genetically modified OA models, such as STR/Ort mice, can broaden the scope of our formulation(*83, 84*). Preclinical investigations are also necessary in higher animals that may require additional optimization of dosages and treatment regimens before translation to humans.

In summary, we report the development of rapamycin as an MP formulation that can be administered after articular joint injury to prevent OA. This formulation was potent to prevent senescence and induced autophagy in articular chondrocytes, reducing inflammatory markers, thus helped prevent and treat post-traumatic OA in the mice model. The relevance of our findings to human disease was validated using chondrocytes isolated from osteoarthritic patients. These findings provide new insights into therapies for senescence prevention and autophagy activation via sustained release formulation to treat trauma-induced osteoarthritis.

## Materials and methods

### Materials

Poly (D, L-lactic-co-glycolic) acid (PLGA, 50:50) of different molecular weights from 10-15 kDa, 75-85 kDa, 85-100 kDa and 190-240 kDa (85:15) with carboxylic acid end groups were purchased from Akina (AP041, AP089, AP036) (West Lafayette, Indiana, USA) and Sigma (739979) (St Louis, MO, USA) respectively. Poly (vinyl alcohol) (PVA, 87~89% hydrolysed, Mw 13,000~23,000kDa) was purchased from Sigma (363170) (St Louis, MO, USA). Rapamycin (or Sirolimus, 99.5% purity) was procured from Alfa *Ae*sar (J62473) (Ward Hill, Massachusetts, USA). Insulin/ transferrin /selenium (ITS) purchased from Gibco (Carlsbad, California, USA) were used for micromass culture of human chondrocyte cell line (C28/I2). All other chemicals were procured from Sigma Aldrich and Thermo Scientific as analytical grade and used as received. Surgical sutures (poly (glycolic acid) based absorbable sutures with ¾ reverse cutting edge needle) were obtained from Dolphin sutures.

#### Antibodies used

Rabbit Anti MMP-13 polyclonal antibody (Abcam, Cambridge, UK, ab39012), Rabbit Anti ADAMTS-5 polyclonal antibody (Abcam, Cambridge, UK, ab41037), Rabbit Anti p16^INK4a^ polyclonal antibody (Thermo Fischer, Waltham, Massachusetts, USA, PA1-30670), Rabbit Anti LC-3B polyclonal antibody (Thermo Fischer, Waltham, Massachusetts, USA, PA1-16930), Alexa Fluor 488 goat anti-rabbit IgG (H+L) (Abcam, Cambridge, UK A11008), HRP conjugated goat anti-rabbit secondary antibody (Thermo Fischer, Waltham, Massachusetts, USA, 31460).

#### Cell line and culture conditions

The human immortal chondrocyte cell line – C28/I2 (Merck) was used in this study. Cells were cultured in DMEM/F-12 (Ham) media (Invitrogen) containing 1 mM sodium pyruvate, 10 mM of HEPES, 140 mM of glucose. The media was supplemented with 10% fetal bovine serum (US origin, Sigma) and 1% antibiotic cocktail containing penicillin/streptomycin (Invitrogen) in an atmosphere of 5% CO_2_ and 37° C.

#### Mice

All the animal experiments were approved by the Institutional Animal Ethics committee (IAEC) (CAF/Ethics/612/2018 and CAF/Ethics/808/2020). IAEC guidelines were followed for the design, experimentation, and analysis of animal experiments. Mice were housed at the Central Animal Facility, Indian Institute of Science (IISc) in individually ventilated cages under monitored temperature and humidity with automated 12 h light-dark cycles. Mice were fed with standard laboratory food and water. Female wild-type (WT) mice of C57BL/6 strain were used for intra-articular injection studies and male wild-type (WT) mice of C57BL/6 strain were used for OA studies.

#### Human cartilage samples

All the human *ex-vivo* experiments were approved by the Institutional Human Ethics committee (IHEC) (02/31.03.2020) and M.S. Ramaiah Medical College, Ethics Committee (MSRMC/EC/AP-06/01-2021). IHEC and MSRMC/EC guidelines were followed for the design, experimentation, and analysis of human experiments. Human knee joint surfaces from femur and tibia were obtained from patients undergoing total knee replacement due to end stage OA, from Ramaiah Medical College and Hospital.

#### Sample’s collection consent statement

The human articular cartilage samples collected to perform this study were obtained from patients with OA undergoing total knee arthroplasty from M.S. Ramaiah Medical College according to IISc and MSRMC approved ethical protocol. Patients in the registry at MSRMC gave informed consent to donate knee joints for research. The information about the study and informed consent sheet was provided to the participants in the research. The information obtained by the analysis of the samples was treated under the criteria of confidentiality at all times.

### Methods

#### Synthesis of PLGA microparticles

PLGA microparticles were synthesized from PLGA polymer of various molecular weights, using the single emulsion technique as described earlier(*38*). Briefly, 100 mg of PLGA was dissolved in 2 mL of dichloromethane (DCM) with or without rapamycin (1 mg), and the homogenization was carried out in 1% polyvinyl alcohol (10 mL) at 12,000 rpm. This solution was added to 1% PVA (110 mL) and was allowed to stir continuously for 3–4 h to evaporate DCM completely. The solution was then centrifuged at 11,000x *g* and the pellet was washed using deionized water twice to wash away the excess PVA. The microparticles were then re-suspended in deionized water and rapidly frozen at −80 °C followed by lyophilization. The lyophilised particles were used for *in vitro* and *in vivo* studies by making a suspension in 1x PBS according to required concentrations and sterilising them under UV for 20 mins. For determining the size of PLGA microparticles, 50 μg particles were dispersed in deionized water (1 mL) and sonicated briefly before analysis. Particle size distributions were determined using dynamic laser light scattering (Nano ZS ZetaSizer (Malvern Instruments, Worcestershire, UK)).

#### Physiochemical characterisation of the microparticle formulation

##### Encapsulation efficiency (EE)

To estimate the amount of rapamycin that was encapsulated in PLGA particles, a standard curve was first prepared by adding different concentrations of rapamycin in DMSO solution containing PLGA (10 mg mL^−1^). The absorbance was read using SPL UV Max™ plate (33096) at 278 nm using plate reader (Tecan™ Microplate Spectrophotometer). The rapamycin-PLGA microparticles (10 mg mL^−1^) were weighed and dissolved in DMSO and the amount of rapamycin encapsulated was determined from the standard curve. EE was calculated using the following equation:

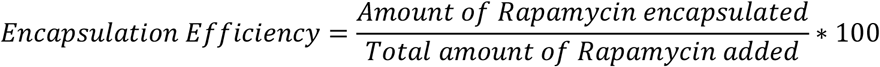

##### *In vitro* release profile of rapamycin/Cy7 loaded microparticles

Rapamycin or Cy7 loaded microparticles of different molecular weights (1 mg) were suspended in 1x PBS (1 mL) solution (pH 7.4) at 37° C in a rotatory spinner. At different incubation times, the particles were pelleted at 11,000x *g* and dissolved in 100 μL DMSO. The dissolved particles were quantified through plate reader instrument (Tecan™ Microplate Spectrophotometer) with pre-determined parameters set to record the absorbance at 278 nm for rapamycin and fluorescence intensity at 750/773 nm for Cy7 dye. The amount of rapamycin/ Cy7 released at each time point was determined from the standard curve made using PLGA spiked with rapamycin/Cy7. The experiment was carried out in triplicates along with blank microparticles as controls.

##### Isolation of human articular chondrocytes from human knee joint surfaces

Knee joints excised during surgery were immediately placed in sterile containers with 1x PBS and brought to the tissue culture facility within an hour. The obtained human articular knee joint surfaces were briefly rinsed in 1x PBS (containing 1% penicillin-streptomycin) once before scraping the cartilage from the underlying bone surface into thin slices of 1-3 mm^3^. These slices were placed in a 10 cm dish and were washed 3 times with 1x PBS. The slices were then incubated at 37° C for 30 minutes in DMEM-F12 media containing pronase (1mg mL^1^). Slices were again rinsed with 1x PBS and were incubated at 37° C for 16-18 h in DMEM-F12 media containing collagenase A solution (3mg mL^−1^). After 16 – 18 h, the clumps were broken apart using a pipette and the cells were pelleted down at 1000x *g* for 10 minutes. The cells were washed with 1x PBS thrice. Finally, the cells were resuspended in DMEM-F12 media, counted, and seeded at a density of 1×10^6^ cells mL^−1^. The media was changed every two days and the cells were subsequently sub-cultured and used for experiments within the first two passages.

##### Senescence induction assay

Primary human articular chondrocytes (HACs) were plated in a 24 well plate with 15,000 cells per well in the untreated group and 30,000 cells per well for other treatment groups. The chondrocytes (C28/I2) were plated in a 24 well plate with 7,500 cells per well in the untreated group and 15,000 cells per well for other treatment groups to maintain cells under sub confluency until the end of the experiment. They were subjected to different stress conditions such as genotoxic stress agent- BrdU (200 μM) or oxidative stress agent- hydrogen peroxide (H_2_O_2_) (200 μM) to simulate inflammation in post-traumatic OA. The primary treatment groups were, vehicle treated cells (DMSO), BrdU/ H_2_O_2_ treated cells, BrdU/ H_2_O_2_ along with free rapamycin (1 μM) co-treated cells, BrdU/ H_2_O_2_ along with RMPs (equivalent to 1 μM) co-treated cells, BrdU/ H_2_O_2_ along with BMPs co-treated cells. Colorimetric SA-β Gal activity was used to stain for senescent cells as previously described (*85*). After exposing the cells to these stress agents for 48 h, cells were washed thrice with 1x PBS and then fixed with fixative solution containing 2% formaldehyde and 0.2% glutaraldehyde in 1x PBS for 10 mins. Following fixation, cells were incubated in SA-β Gal staining solution (1 mg mL^−1^ 5- bromo-4-chloro-3- indolyl-beta-d-galactopyranoside (X-Gal), 1x citric acid/sodium phosphate buffer (pH 6.0), 5 mM potassium ferricyanide, 5 mM potassium ferrocyanide, 150 mM sodium chloride, and 2 mM magnesium chloride) at 37° C overnight. The enzymatic reaction was stopped after 16 h and cells were washed three times with 1x PBS. Five random bright field images were taken per well. These images were analysed using ImageJ software.

To automatically score the senescent cells, we developed a custom-built macros algorithm to score the senescent cells based on their size and the intensity of SA-β Gal staining. Bright field images were taken from each treatment groups (vehicle (1x PBS) treated cells, H_2_O_2_ treated cells, H_2_O_2_ and rapamycin/RMPs treated cells etc). The macros algorithm was run to count the total number of senescent cells.

##### RMPs treatment in senescence induction assays (Micromass culture)

We generated micromasses as described previously(*55*). Briefly, the cells were seeded as a 15 μL suspension in growth media in a 24 well plate at a density of 2.5 × 10^7^ cells mL^−1^. The cells were allowed to adhere to the well plate for 3 h after which growth media was added. After 24 h, the growth media was changed to differentiation media containing supplements (Insulin/Transferrin/Selenium, TGF-β (10 ng mL^−1^), and ascorbic acid). Micromasses were treated with BrdU (600 μM) /H_2_O_2_ (100 μM) and co-treated with free rapamycin (1 μM)/RMPs (final concentration 1 μM). After 48 h of incubation, the micro masses were fixed using 4% formaldehyde followed by Alcian blue staining at pH<1 to stain the sGAG overnight. After 16 - 18 h, micro masses were washed with deionised water to remove any non-specific stains followed by Alcian blue stain extraction using 6M Guanidine HCl. The extracted Guanidine HCl’s absorbance was read at 630 nm using a plate reader (Tecan™ Microplate Spectrophotometer) to quantify the sGAG present in the micro masses after various treatments. Similar experiments were carried out with incubation time of up to 8 days with BrdU/H_2_O_2_ and BrdU/H_2_O_2_ along with free rapamycin/RMPs for 8 days. Media was changed every two days and were replenished with BrdU/H_2_O_2_ along with respective treatments.

##### Immunocytochemistry

After 48 h of treatment, the cells were fixed using 4% PFA followed by permeabilization using 0.05% Triton X-100. The cells were then blocked using 5% non-fat dry milk and incubated overnight at 4° C with rabbit anti LC-3B polyclonal antibody (0.5 μg mL^−1^). After 16 – 18 h, the cells were washed and incubated with Alexa Fluor 488 goat anti-rabbit IgG (H+L) (1 μg mL^− 1^) for 1 h followed by nuclear staining using DAPI (300 nM) for 10 mins. These cells were then visualised using FITC (495/519 nm) and DAPI (358/451 nm) channel using IN Cell Analyzer 6000 (GE Life Sciences) at 40x magnification.

The obtained images were quantified for the LC3B puncta per cell using a custom-built automated macros algorithm. For counting LC3B puncta, FITC channel images threshold was adjusted so that the individual puncta are picked up and counted. The same threshold values were used for analysis of all the images in the experiment. Similarly, the total cell numbers were obtained by an automated macro algorithm picking up the DAPI stained nucleus to give the total cell number. The average puncta per cell were obtained by dividing the total puncta per image by the total number of cells in that image.

##### Residence time of PLGA MPs in mouse knee joint

PLGA microparticles of molecular weight 75 kDa - 85 kDa with Cy7 dye were suspended in 1xPBS at a concentration of 20 mg mL^−1^. Ten μL of this formulation was injected in mice left knee joint by intra-articular injections on Day 0. The contralateral legs were used as control and were given equivalent dose of free Cy7 dye. The mice (n=5 per group) were anesthetized by isoflurane prior to transfer in the imaging system (IVIS^®^ Spectrum In Vivo Imaging System), and fluorescence signal was measured with predetermined exposure time. Imaging (fluorescence) was performed on Day 0 (pre- and post-injection), 1, 3, 5, 7, 14, 21, 30 and 35 using the IVIS. Radiance efficiency (p s^−1^ sr^−1^ μW^−1^) within a region of interest (ROI) was quantified by the IVIS^®^ Spectrum Living imaging software.

##### Effect of RMPs on Surgically Induced Osteoarthritis

Male C57BL/6 mice (25g) were used for this study(*86*). Following an intraperitoneal injection of ketamine (80 mg kg^−1^) and xylazine (10 mg kg^−1^) as a cocktail in sterile 1x PBS, the animals were rested until the surgical plane was reached with no active plantar reflexes. The knee joint capsule was exposed by medial parapatellar incision. After the joint capsule was opened, the patella was displaced laterally followed by transection of the medial menisco tibial ligament using surgical blade number 11 and scalpel. Following irrigation of the operated site with saline, the capsule and skin were sutured separately using absorbable PGA sutures. Sham groups received only an incision to expose the medial collateral ligament which was then sutured back like the other DMM operated groups. Postoperatively, the mice were not immobilised and were allowed to move freely in the cage. Each experimental group was evaluated by gross morphological examination for swelling, pain or change in gait of the animal post-surgery and during the entire duration of the experiment.

#### Classification of study

##### Prophylactic dose study

Group 1- DMM operated animals with no treatment (4 mice)

Group 2- Surgical control receiving no treatment (Sham) (4 mice)

Group 3- DMM operated with Free rapamycin injection (1.8 μg) (6 mice)

Group 4- DMM operated with Free rapamycin injection (180 ng) (5 mice)

Group 5- DMM operated with RMP injection (200 μg particles containing equivalent of 1.8 μg rapamycin) (6 mice)

Group 6- DMM operated with RMP injection (200 μg particles containing equivalent of 180 ng rapamycin) (6 mice)

Group 7- DMM operated with BMP injection (200 μg particles) (4 mice)

##### Therapeutic dose study

Group 1-DMM operated animals with no treatment (4 mice)

Group 2- Surgical control receiving no treatment (Sham) (4 mice)

Group 3- DMM operated with Free rapamycin injection (1.8 μg) (4 mice)

Group 4- DMM operated with RMP injection (200 μg particles containing equivalent of 1.8 μg rapamycin) (6 mice)

Group 5- DMM operated with BMP injection (200 μg particles) (4 mice)

In prophylactic study, IA injection was given 7 days after DMM surgery, followed by injections at day 24 and day 42 with euthanasia at day 60. In therapeutic study, injection was given at 3 weeks after DMM surgery, followed by injections at week 6 with euthanasia at week 9.

##### Histology

The whole joints of operated knees were fixed in 4% paraformaldehyde for 24 h. After decalcification in 5% formic acid for five days, the joints were embedded in paraffin. Five-micron sections were stained with safranin-O-fast green staining using the described protocol(*87*). In brief, slides were deparaffinized with xylene and alcohol. After a 5-min wash in running tap water, slides were stained with fresh Wiegert’s iron haematoxylin for 10 min. Following a 5-min wash in running tap water, slides were soaked in 0.1% safranin-O (10 min) and 0.05% fast green (5 min) in this order. After dehydration with alcohol and xylene, the slides were stained with DAPI. The prepared slides were graded according to the OARSI osteoarthritis cartilage histopathology assessment system (*88*). The examination was performed by veterinarian in a blinded manner to minimize observer bias.

##### Immunohistochemical studies

Sections were processed similar to Safranin O staining until the alcohol step followed by heat induced epitope retrieval at 95° C in 1x TBST buffer (pH 9), followed by blocking using blotto (0.8%). The slides were rinsed in 1x TBST and incubated with MMP-13, ADAMTS-5, LC3B and p16^INK4a^ antibodies (1:300) diluted in blocking solution at 4° C for 16 – 18 h. The slides were again washed in 1x TBST followed by incubation with HRP conjugated secondary antibody (1:150) diluted in blocking solution in room temperature for 2 h. The slides were again rinsed and DAB (3,3’-Diaminobenzidine) substrate was added and incubated for 1 h. The slides were then washed, dried and cover slipped using mounting media before brightfield imaging. The number of cells positive in each image was quantified using an automated macros program in ImageJ software.

##### Statistical analysis

All statistical analysis was performed using Graph Pad Prism 6. Data were presented as mean ± standard deviation. Non-linear regression (least square method) two phase exponential decay curve was used to fit rapamycin release profile with constrains of maximum value as 100 and minimum value as 0. Differences between groups were analysed by t-test or one-way analysis of variance (ANOVA) and non-parametric data was analysed using Mann Whitney U test or Kruskal Wallis test with Tukey’s multiple comparison test, with *p* < 0.05 considered significant. Since the estimate for the mean and standard deviation for each of the animal experimental group was not available initially, we kept a minimum of 4 animals in each group. After the results were available, we retrospectively calculated the statistical power to detect differences between free drug and RMPs using G*Power 3.1 software. The power for both prophylactic and therapeutic mice OA studies was greater than or equal to 80%.

## Acknowledgements

We thank Central Animal Facility (CAF) for breeding and maintaining mice and access to IVIS facilities which was facilitated by Ms Lekha Kandasami. Dr. Rakshith Kumar and Dr. Rekha Balu performed the blinded OARSI scoring.

## Funding statement

We thank Mr. Lakshmi Narayanan, Early Career Research Award (Science and Engineering Research Board, Department of Science and Technology, India, ECR/2017/002178), Har Gobind Khorana Innovative Young Biotechnologist Award (Department of Biotechnology, India, BT/12/IYBA/2019/04), and the Indian Institute of Science, Bangalore start-up for funding our project. Funding from Dr Vijaya and Rajagopal Rao for Biomedical Engineering research at the Centre for BioSystems Science and Engineering is also acknowledged.

## Author contributions

Study design and experimental planning: KMD and RA, Surgical assistance: AAD and SA, Chief surgeons providing human samples: RKS, AKP, Supervision: RA, Funding acquisition: RA, Writing – original draft: KMD, Writing – review & editing: KMD, AAD, SA and RA. All authors approved the final version to be published.

## Competing interests

The other authors declare that they have no competing interests.

## Notes

### Competing Interest Statement

The authors have declared no competing interest.

