## Supplementary Information for "Rapamycin microparticles induce autophagy, prevent senescence and are effective in treatment of Osteoarthritis"

*Author's affiliations:*

[1]*Centre for BioSystems Science and Engineering, Indian Institute of Science, Bengaluru, India 560012.*

[2]*Department of Orthopedics, M.S. Ramaiah Medical College, Bengaluru, India 560054.*

**Supplementary information**

Fig. S1. Representative human tibial articular surface bearing lesions from OA patients.

Fig. S2. Rapamycin MPs induce autophagy in chondrocytes.

Fig. S3. Rapamycin MPs prevent senescence in chondrocytes (C28/I2) under genotoxic stress.

Fig. S4. Rapamycin MPs prevent senescence in chondrocytes (C28/I2) under oxidative stress.

Fig. S5. Rapamycin MPs prevent loss of sGAG in micromass cultures exposed to genotoxic and oxidative stresses.

Fig. S6. I*n vitro* Cy7 release curve follows a similar trend of rapamycin release from PLGA MPs.

Fig. S7. PLGA MPs exhibit prolonged residence time in mice knee joints.

Fig. S8. RMPs in prophylactic regimen reduced the markers of inflammation associated with OA in surgically induced murine OA models.

Fig. S9. RMPs in therapeutic regimen reduced the markers of inflammation associated with OA in surgically induced murine OA models.

Table S1: PLGA microparticles of different molecular weights and PLA: PGA ratios, their respective sizes, and their rapamycin encapsulation efficiency.

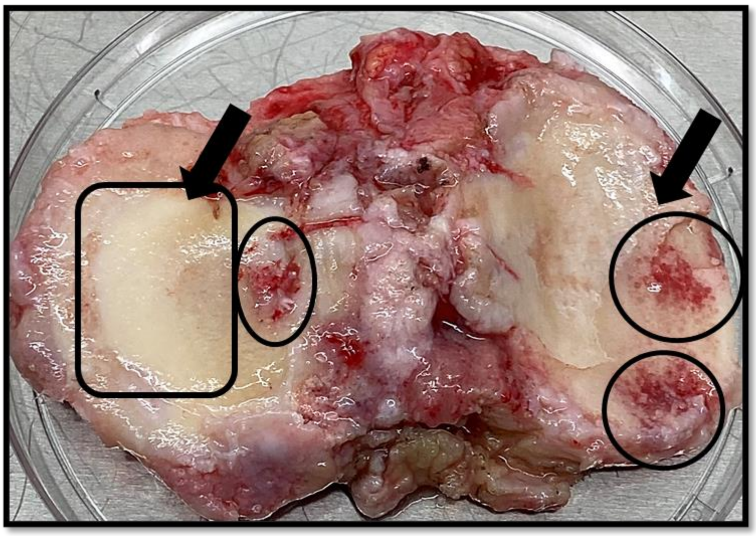

**Fig. S1. Representative human tibial articular surface bearing lesions from OA patients.** The circles represent the areas bearing OA lesions with involvement of subchondral bone and the rectangular area represents the non-lesioned areas which were used for IHC.

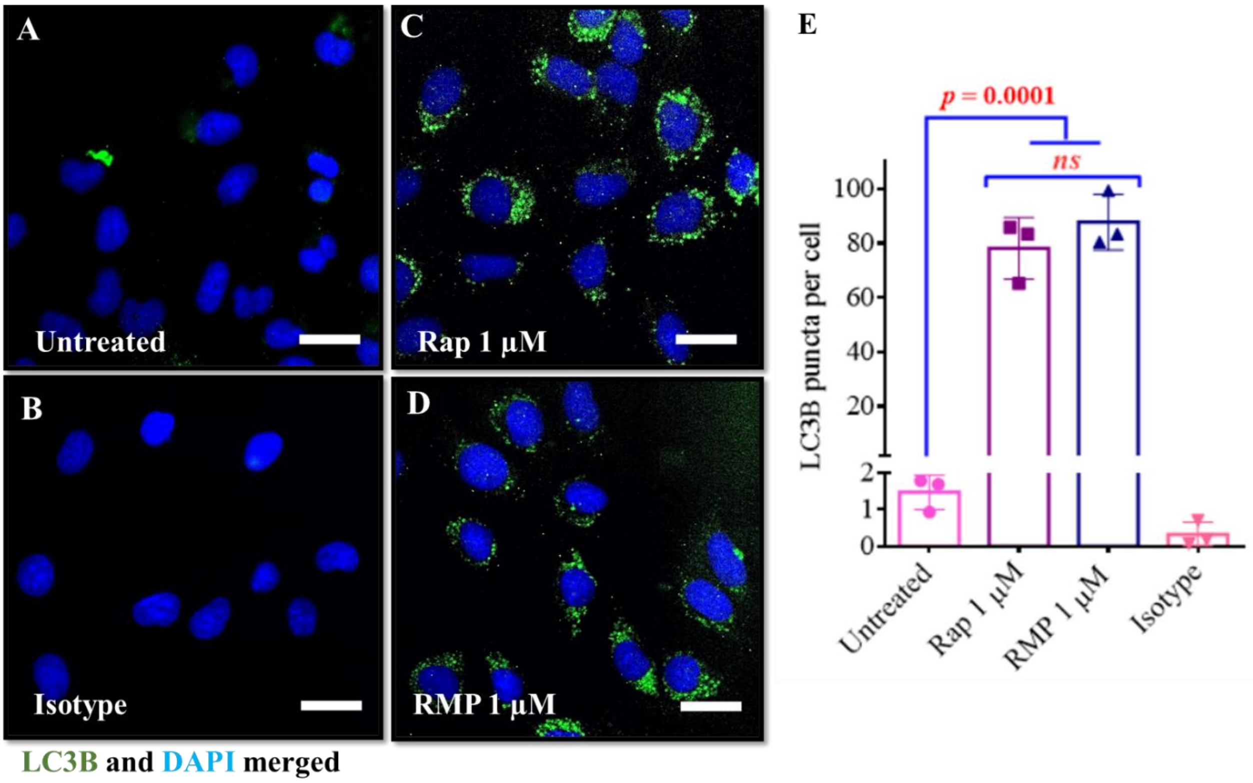


**Fig. S2. Rapamycin MPs induce autophagy in chondrocytes.** Fluorescence microscopy images of C28/I2 cells stained with DAPI and LC3B after (**A**) **no treatment**, **(B)** isotype antibody, **(C)** free rapamycin (1 µM) and **(D)** Rapamycin MPs (1 µM rapamycin) treatment. **(E)** Quantification of LC3B puncta per cell using ImageJ software (n = 3 per group). Data in graphs represent the mean ± s.d. and *p* values were determined by one-way analysis of variance (ANOVA) and Tukey's post hoc tests. *p*-value < 0.05 was considered significant. *ns* - non significant; Scale bar, 20 µm.

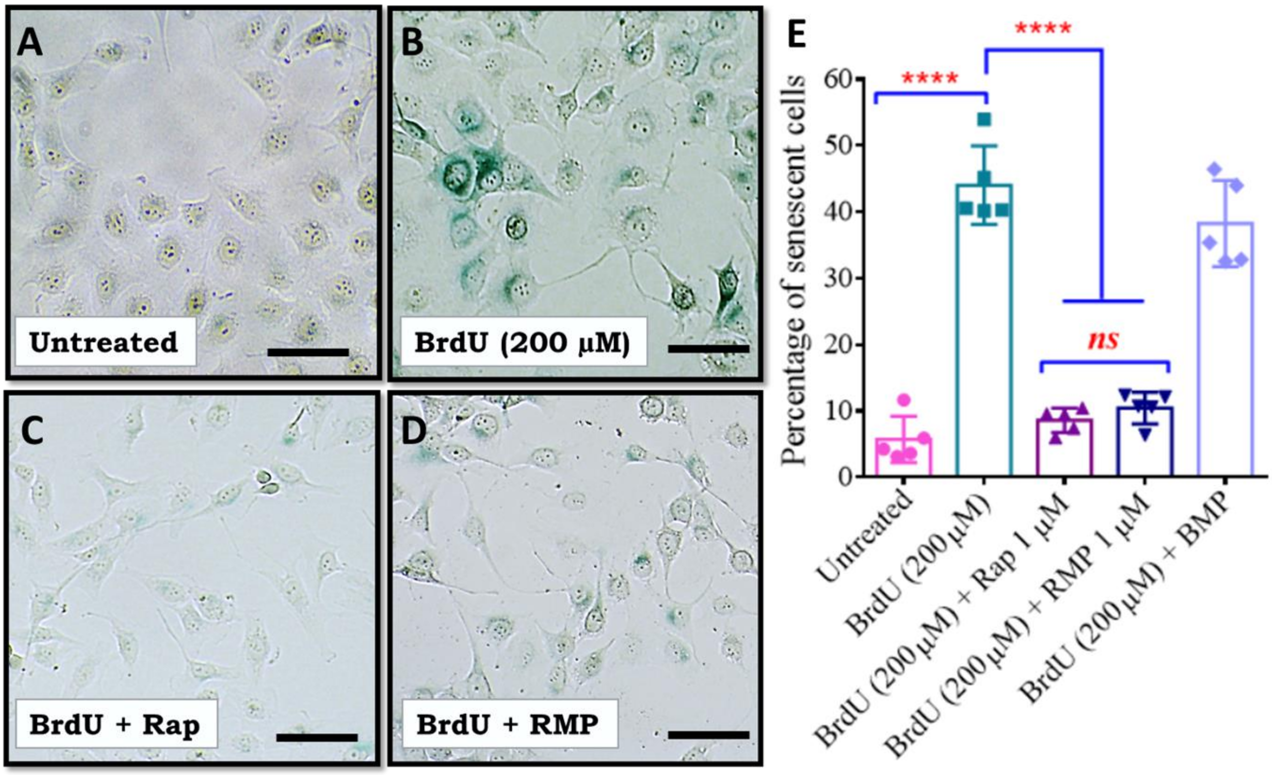


**Fig. S3. Rapamycin MPs prevent senescence in chondrocytes (C28/I2) under genotoxic stress.** SA-β Gal-stained images of C28/I2 cells exposed to **(A) no treatment**, **(B)** genotoxic (BrdU) stress, **(C)** genotoxic (BrdU) stress along with free rapamycin (1µM) treatment and, **(D)** genotoxic (BrdU) stress along with rapamycin MPs (1µM) treatment. **(E)** Percentage of senescent cells under genotoxic (BrdU) stress condition, (n = 5 per group). Data in the graph represent the mean ± s.d. and *p* values were determined by one-way analysis of variance (ANOVA) and Tukey's post hoc tests. *p*-value < 0.05 was considered significant. BMP-Blank Microparticles, RMP- Rapamycin loaded Microparticles. ****$p$<0.0001, *ns* - non significant; Scale bar, 40 µm.

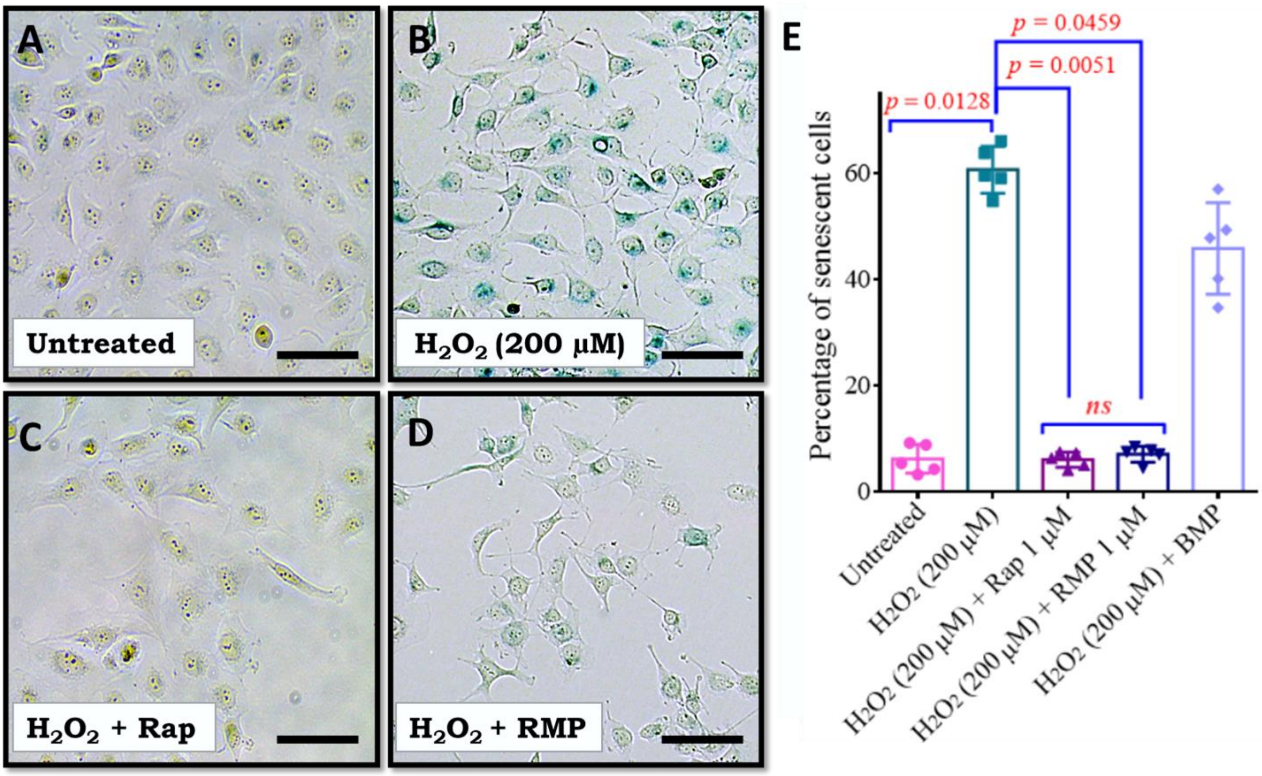


**Fig. S4. Rapamycin MPs prevent senescence in chondrocytes (C28/I2) under oxidative stress.** SA-β Gal-stained images of C28/I2 cells exposed to **(A) no treatment**, **(B)** oxidative ($H_2O_2$) stress, **(C)** oxidative ($H_2O_2$) stress along with free rapamycin (1µM) treatment and, **(D)** oxidative ($H_2O_2$) stress along with rapamycin MPs (1µM) treatment. **(E)** Percentage of senescent cells under oxidative ($H_2O_2$) stress condition (n = 5 per group). Data in the graph represent the mean ± s.d. and *p* values were determined by one-way analysis of variance (ANOVA) and Tukey's post hoc tests. *p*-value < 0.05 was considered significant. BMP-Blank Microparticles, RMP- Rapamycin loaded Microparticles. *ns* - non significant; Scale bar, 40 µm.

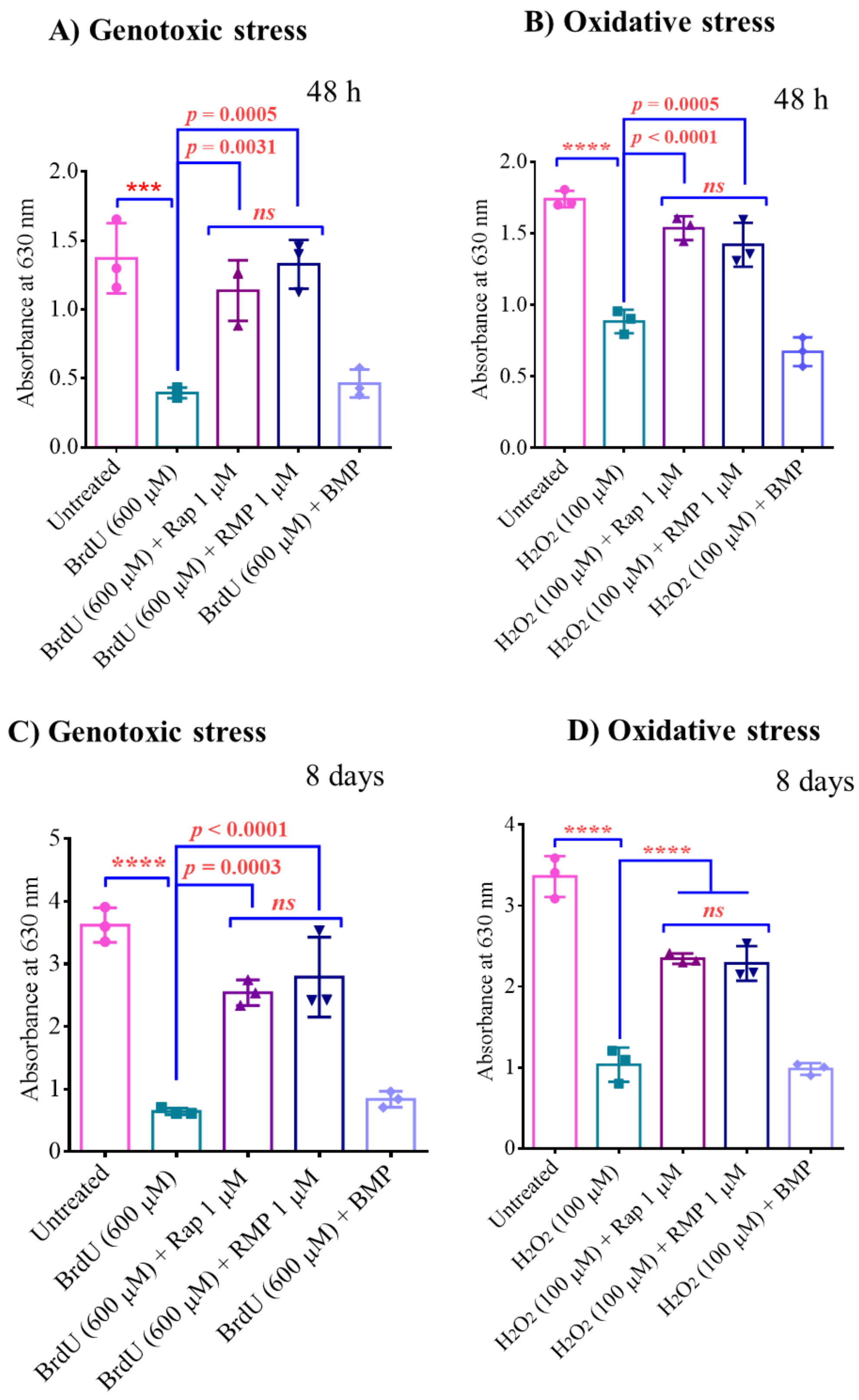


**Fig. S5. Rapamycin MPs prevent loss of sGAG in micromass cultures exposed to genotoxic and oxidative stresses.** sGAG production from C28/I2 micromass culture after treatment with various particle and drug formulations under **(A)** genotoxic (BrdU) stress for 48 h (n = 3), **(B)** oxidative ($H_2O_2$) stress for 48 h (n = 3), **(C)** genotoxic (BrdU) stress for 8 days (n = 3), and **(D)** oxidative ($H_2O_2$) stress for 8 days (n = 3). Data in graphs represent the mean ± s.d. and $p$ values were determined by one-way analysis of variance (ANOVA) and Tukey's post hoc. $p$-value < 0.05 was considered significant. BMP - Blank Microparticles, RMP - Rapamycin loaded Microparticles. ****$p$<0.0001, ***$p$<0.001, *ns* - non significant.

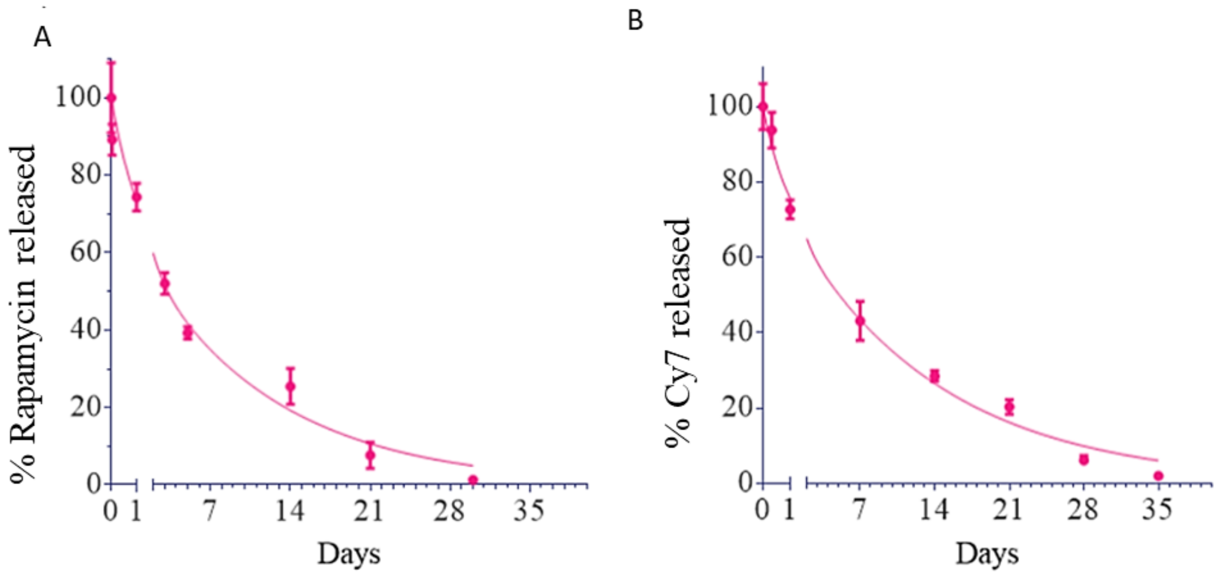


**Fig. S6. *In vitro* Cy7 and rapamycin release curve follows a similar trend from PLGA MPs.** Quantification of *in vitro* release profiles of **(A)** rapamycin and **(B)** Cy7 dye from PLGA particles synthesized from PLGA 75-85 kDa (50:50). Nonlinear regression (least square method) was used to fit a two-phase exponential decay curve in all the release profiles. Data in graphs represent the mean ± s.d. (n = 3).

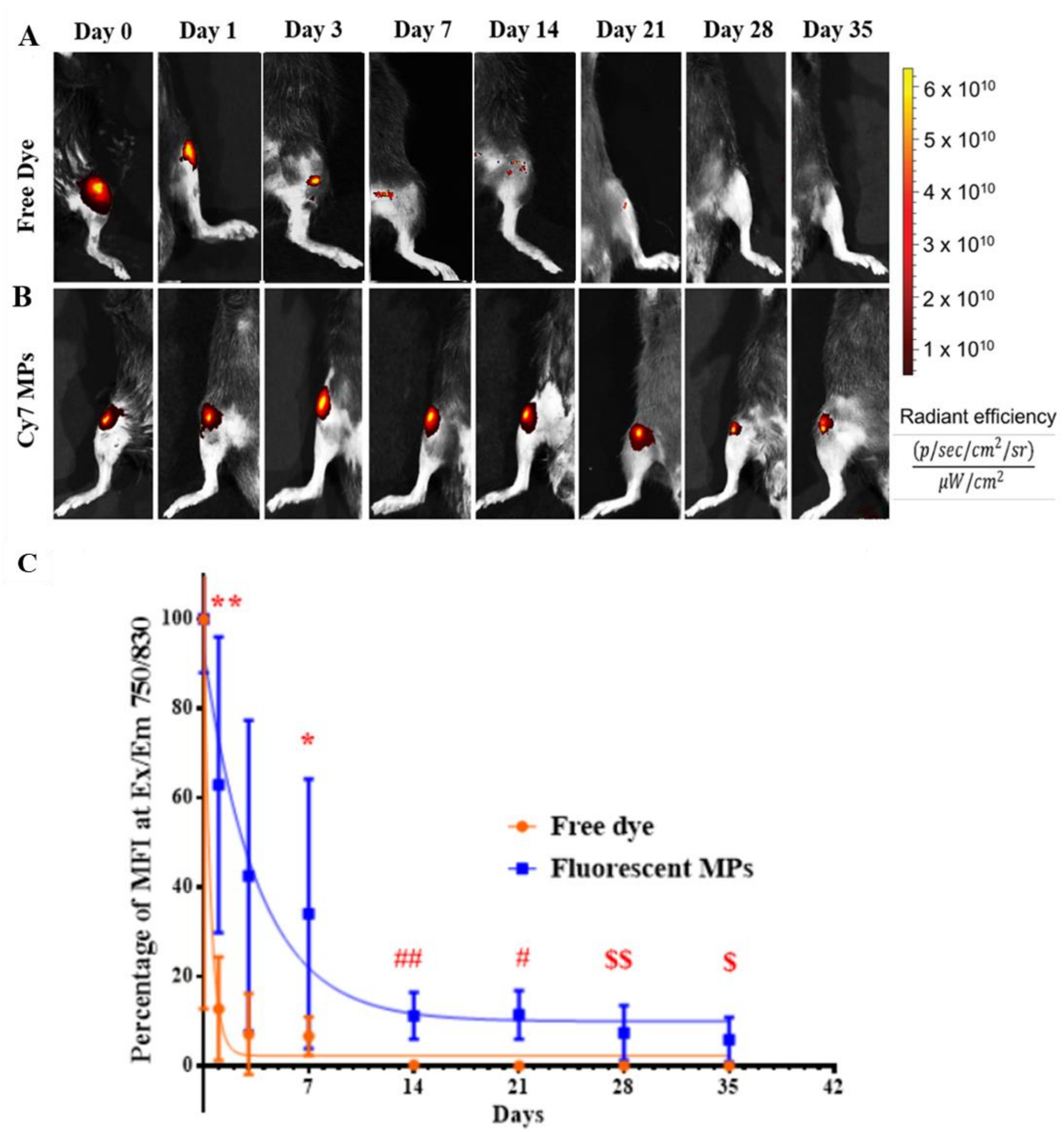


**Fig. S7. PLGA MPs exhibit prolonged residence time in mice knee joints.** *In vivo* images of mice knee joints imaged at0, 1, 3, 7, 14, 21, 28, and 35 days after intra-articular injection of **(A)** free Cy7 dye and **(B)** Cy7-labeled PLGA MPs. **(C)** Percentage fluorescent intensity remaining of injected formulations with respect to days (n = 5 mice per group). Data in graph were fitted with non-linear regression (least square method) one-phase exponential decay curve. Data in graph represent the mean ± s.d. (n = 5 at each time point) and *p* values were determined by unpaired t test or Mann Whitney test. *p*-value < 0.05 was considered significant. ***** - 0.0159, ******- 0.0126, **#** - 0.0016, **##** - 0.0079, **$** - 0.0296, **$$** - 0.0269.

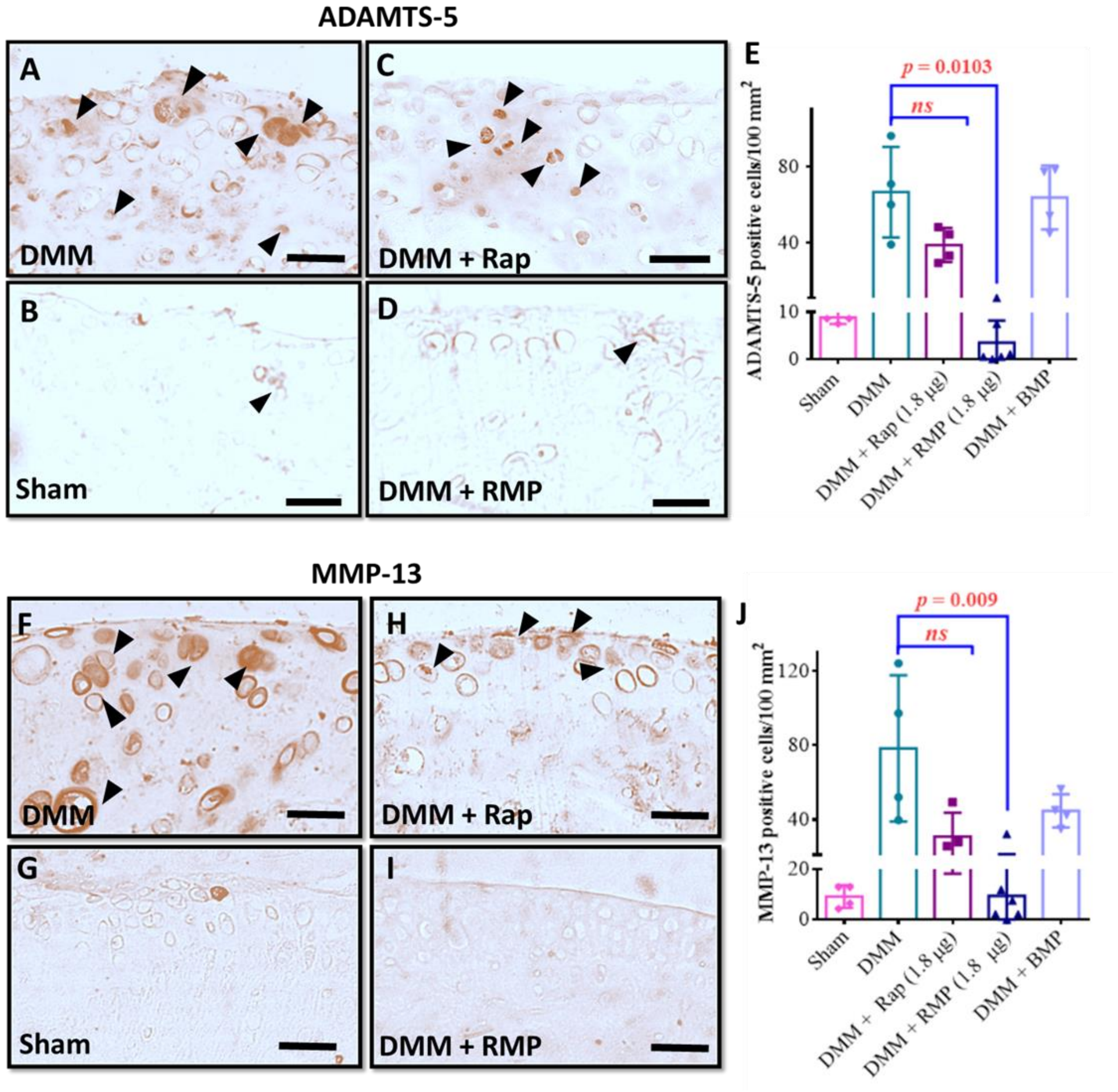


**Fig. S8. RMPs in prophylactic regimen reduced the markers of inflammation associated with OA in surgically induced murine OA models.** Immunohistochemical staining for ADAMTS-5 in **(A)** DMM, **(B)** Sham, **(C)** DMM operated with free rapamycin (1.8 µg) treatment and, **(D)** DMM operated with RMP (200 µg particles containing 1.8 µg rapamycin) treatment. **(E)** Quantification of ADAMTS-5 positive cells per 100 $mm^2$. Immunohistochemical staining for MMP-13 in **(F)** DMM, **(G)** Sham, **(H)** DMM operated with rapamycin (1.8 µg) treatment and, **(I)** DMM operated with RMP (200 µg particles containing 1.8 µg rapamycin) treatment. **(J)** Quantification of MMP-13 positive cells per 100 $mm^2$. The data in graphs represent the mean ± s.d. and *p* values were determined using one-way analysis of variance (ANOVA) or Kruskal Wallis test and Tukey's post hoc analysis. DMM-Destabilization of Medial Meniscus model, BMP - Blank Microparticles, and RMP – Rapamycin Microparticles. *ns* - non significant; Scale bar, 50µm.

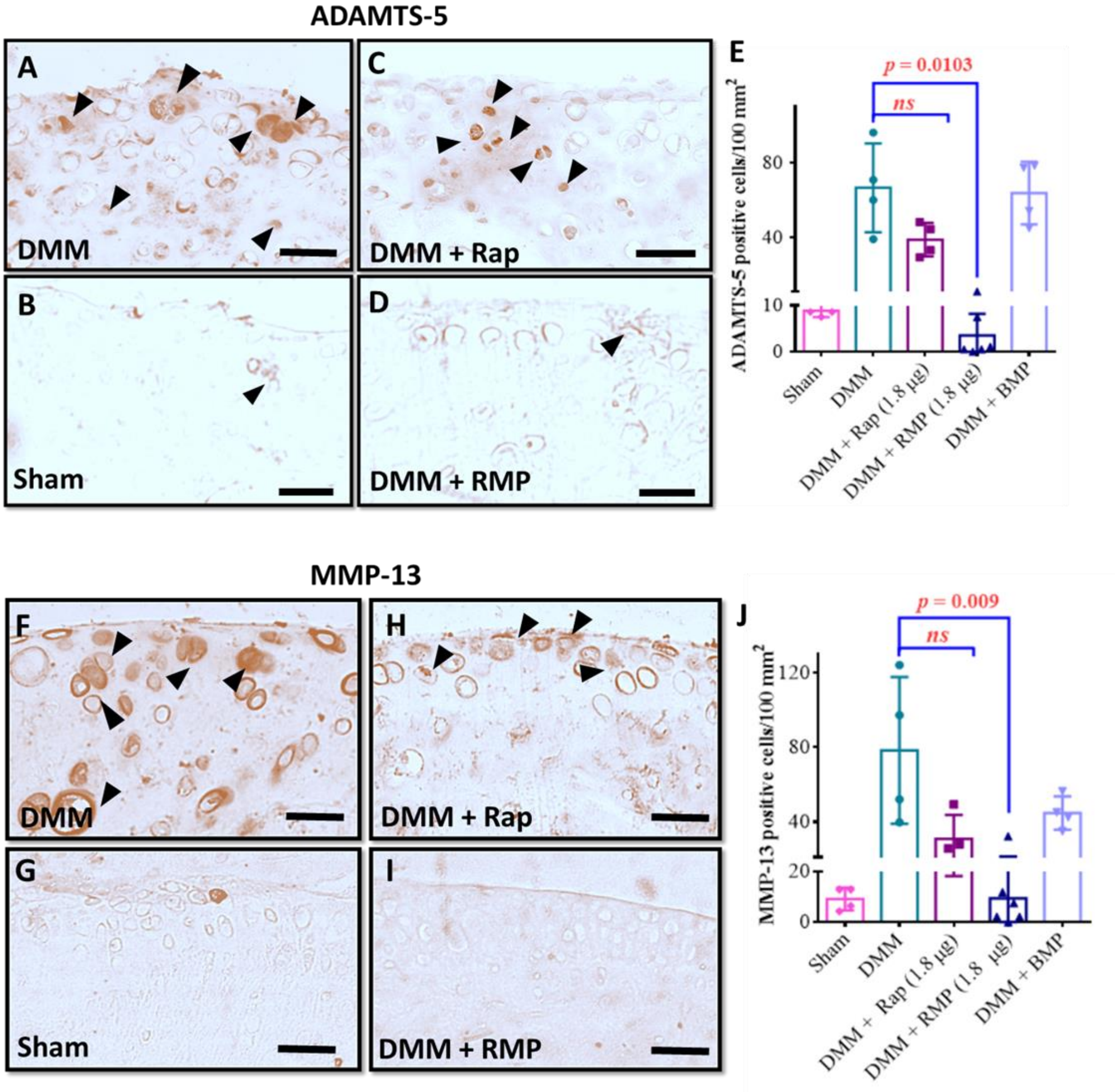


**Fig. S9. RMPs in therapeutic regimen reduced the markers of inflammation associated with OA in surgically induced murine OA models.** Immunohistochemical staining for ADAMTS-5 in **(A)** DMM, **(B)** Sham, **(C)** DMM operated with rapamycin (1.8 µg) treatment and, **(D)** DMM operated with RMP (200 µg particles containing 1.8 µg rapamycin) treatment. **(E)** Quantification of ADAMTS-5 positive cells per 100 $mm^2$. Immunohistochemical staining for MMP-13 in **(F)** DMM, **(G)** Sham, **(H)** DMM operated with rapamycin (1.8 µg) treatment and, **(I)** DMM operated with RMP (200 µg particles containing 1.8 µg rapamycin) treatment. **(J)** Quantification of MMP-13 positive cells per 100 $mm^2$. The data in graphs represent the mean ± s.d. and *p* values were determined using one-way analysis of variance (ANOVA) or Kruskal Wallis test and Tukey's post hoc analysis. DMM-Destabilization of Medial Meniscus model, BMP - Blank Microparticles, and RMP – Rapamycin Microparticles. *ns* - non significant; Scale bar, 50µm.

Table S1: PLGA microparticles of different molecular weights and PLA: PGA ratios, their respective sizes, and their rapamycin encapsulation efficiency.

| **Molecular weight of PLGA (Ratio of PLA: PGA)** | **Size measured by DLS [nm]** | **Rapamycin encapsulation efficiency [%]** |
|---|---|---|
| 10 kDa - 15 kDa (50:50) | 1039 ± 188 | 71.83 ± 4.889 |
| 75 kDa - 85 kDa (50:50) | 1153 ± 174.5 | 27.26 ± 2.704 |
| 190 kDa –240 kDa (85:15) | 1192 ± 232.2 | 51.10 ± 1.720 |
